# Direct Generation of Protein Conformational Ensembles via Machine Learning

**DOI:** 10.1101/2022.06.18.496675

**Authors:** Giacomo Janson, Gilberto Valdes-Garcia, Lim Heo, Michael Feig

## Abstract

Dynamics and conformational sampling are essential for linking protein structure to biological function. While challenging to probe experimentally, computer simulations are widely used to describe protein dynamics, but at significant computational costs that continue to limit the systems that can be studied. Here, we demonstrate that machine learning can be trained with simulation data to directly generate physically realistic conformational ensembles of proteins without the need for any sampling and at negligible computational cost. As a proof-of-principle a generative adversarial network based on a transformer architecture with self-attention was trained on coarse-grained simulations of intrinsically disordered peptides. The resulting model, idpGAN, can predict sequence-dependent ensembles for any sequence demonstrating that transferability can be achieved beyond the limited training data. idpGAN was also retrained on atomistic simulation data to show that the approach can be extended in principle to higher-resolution conformational ensemble generation.

## INTRODUCTION

The biological function of a protein is determined not just by a single three-dimensional (3D) structure but its dynamical properties that give rise to conformational ensembles ^1^. Characterizing conformational ensembles is therefore crucial to mechanistically understand the activity of proteins and their regulation, and has an impact on biomedical sciences, biotechnology and drug design ^2-4^.

Experimental techniques for probing the structural dynamics of biomolecules are laborious and suffer from low spatial or temporal resolution ^5^. For this reason, computational methods are often employed to investigate protein dynamics and generate structural ensembles. A powerful strategy in this field is the use of physics-based molecular dynamics (MD) simulations ^6^. In MD, the goal is to sample from the distribution of possible configurations of a molecular system to identify the energetically most favorable regions in conformational space. However, because of high dimensionality and significant kinetic barriers, this presents a formidable computational challenge for all but the very simplest protein systems, even with specialized computer hardware ^7^ or when enhanced sampling methods are applied ^8^. Therefore, alternative strategies for accelerating the generation of biologically relevant dynamic ensembles for a given protein are needed.

In recent years, data-driven machine learning techniques have proven to be extremely fruitful in tackling the protein structure prediction problem ^9^, where the goal is to predict a single 3D conformation of a protein given its amino acid sequence. Predictions from advanced machine learning methods, such as AlphaFold 2 (AF2) ^10,11^, have reached remarkable accuracy in faithfully matching the experimentally-determined ensemble-averaged structures of proteins ^12^. However, since proteins are dynamic entities with multiple conformational states, these methods provide incomplete information. This is especially true for intrinsically disordered proteins (IDPs) ^13^, molecules that lack a stable structural state and exhibit high conformational variability^14,15^.

Given the success of machine learning methods in protein structure prediction and numerous other scientific problems, they are also a promising strategy for accelerating the generation of protein dynamics and conformational ensembles. Currently, multiple strategies for harnessing machine learning models in this field have been explored, including the use of models to facilitate the analysis of complex molecular simulations ^16^, to guide MD sampling ^17^ or to provide optimized energy functions ^18^. Another strategy, and the one followed in this work, is to directly model molecular conformational ensembles through a class of methods called generative models ^19^.

Generative models are based on neural networks and have been proven to be effective in several artificial intelligence tasks ^20,21^. In the context of generating conformational ensembles, such models may be trained on datasets of molecular conformations obtained by “classical” computational methods, such as MD. The idea is essentially to learn the probability distribution of the conformations in a given training set and, once trained, such models can be used to quickly draw statistically-independent samples from these complex, highly-dimensional distributions ^22,23^. Because generative models are not subject to kinetic barriers, they are a powerful strategy for circumventing the computationally expensive sampling via MD.

For generative models to have real utility in substituting for MD simulations, it is essential that previously unseen molecular conformations can be generated for a given system, and that conformations can be generated for new systems with different chemical compositions from what was used in the training set. This can be achieved in principle with conditional generative models that are trained with data from multiple molecules by taking their atomic composition as conditional information. Interestingly, it has been shown that, when trained with sufficiently large datasets, it is possible to generate realistic conformations for molecules unseen in training. This suggests that such models learn not just the probabilities of different conformations encountered in the training set, but that they can learn transferable features of how favorable conformations are constructed. Currently, conditional generative models have been applied successfully only on small molecules ^24-26^. While there have been advances in the unconditional modeling for proteins ^27^, conditional modeling for more complex molecules such as proteins has not been explored yet. Developing an accurate conditional model for proteins would have a dramatic impact on the field of protein dynamics, because it could potentially serve as a direct and computationally efficient generator of conformational ensembles of any protein sequence.

In this study, we present what is to our knowledge the first conditional generative model for protein molecules and apply it to model the conformational ensembles of IDPs. We chose to work with IDPs because of their conformational variability, which we aim to capture using machine learning. Given the complexity of the problem, we work on a simplified description based on a coarse grained (CG) representation. Our training data consists of MD simulations of IDPs obtained using a residue-level CG force field (FF) developed by us. Such CG models capture amino acid-dependent residue interactions in aqueous solvent to match experimental properties of IDPs such as their radius-of-gyration distributions or condensation propensities ^28^. We use a Generative Adversarial Network (GAN) ^29^ to learn the distribution of 3D conformations in the MD data. Our model, which we call idpGAN, has a network architecture that incorporates ideas from machine learning models used in protein structure prediction ^10^. It directly outputs 3D Cartesian coordinates and can model previously unseen conformations of CG proteins of variable sequences and lengths. Since GANs have fast sampling capabilities, our model can generate thousands of independent conformations in fractions of a second, providing a computationally efficient way to reproduce MD conformational ensembles. To show that our method can be adapted to higher resolution protein representations, we also employ it to model the dynamics of α-synuclein as observed in all-atom simulations ^30^. Finally, we discuss the strengths and weaknesses of our approach, and reason over the challenges for generating atomistic conformational ensembles for any protein system.

## RESULTS

### IdpGAN network architecture and training

IdpGAN is a generative model trained on MD data to directly output 3D molecular conformations at a Cα coarse-grained level. From different types of generative models, we chose here GANs ^29^ because of their reported ability to generate high-quality samples and their fast sampling capabilities ^19^. As shown in **Fig. 1**, the learning process of a GAN involves an adversarial game between two neural networks, a generator (G) and a discriminator (D).

**Fig. 1.**
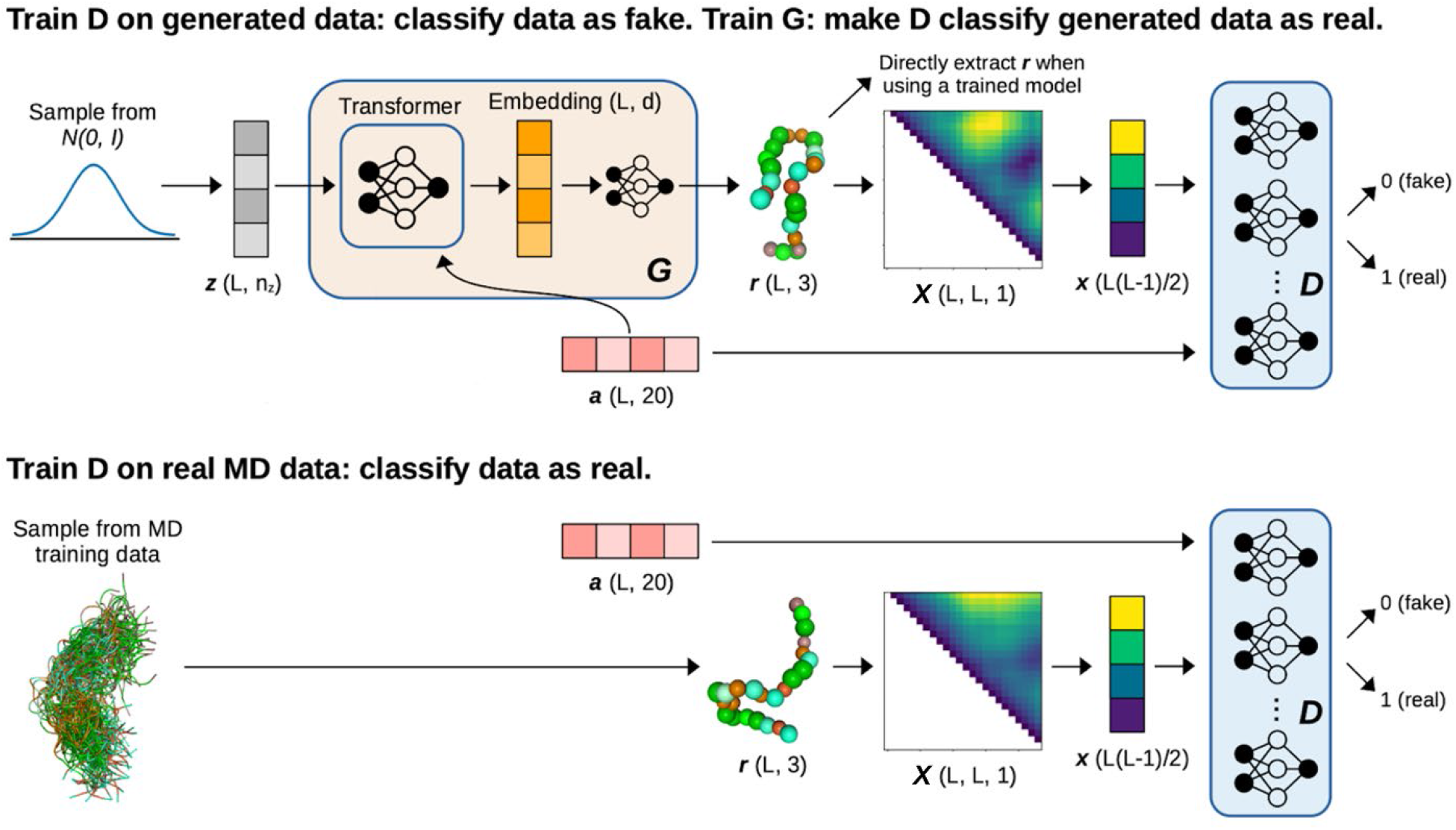
Overview of the idpGAN network architecture. A ***z*** latent sequence is used as input to the G network. Amino acid information ***a*** is also provided as input to G. The output of the last transformer block of G is mapped to 3D Cartesian coordinates ***r*** through a position-wise fully-connected network. Conformations ***r*** from G and the training set are converted in distance matrices and their upper triangles are used as input to a set of D networks, which also receive as input ***a***. The objective of the D networks is to correctly classify real (MD) and fake (generated) samples. The objective of G is to generate increasingly realistic samples to decrease the performance of D. Once idpGAN is trained, the generated coordinates ***r*** from G can be directly used without converting them to distance matrices.

The G network of idpGAN is based on a transformer architecture ^31^. When generating a conformation for a protein with *L* residues, a latent sequence 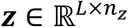 is sampled (its values are randomly extracted from a normal prior). The G network takes as input ***z***, progressively updates it through a series of transformer blocks (we use 8) that produce intermediary embeddings, and finally outputs a sequence ***r*** ∈ ℝ^*L*×3^ corresponding to the 3D coordinates of the Cα atoms of the protein. In addition to ***z***, the network also takes as input a sequence ***a*** ∈ ℝ^*L*×20^ that contains one-hot encoding for amino acid types. This latter data provides the conditional information used to model proteins with different amino acids. Transformer-like architectures are the cornerstone of AF2 ^10^ and their characteristics are well-suited for protein conformation generation. First, they naturally work with variable-size outputs, thus allowing proteins of different lengths to be modeled. Additionally, they use a self-attention mechanism ensuring that each of the *L* tokens in an embedding sequence (that correspond to residue representations) are updated using information from the rest of the sequence, thus helping to form consistent 3D structures ^9^. The “Methods” section and **Supplementary Fig. 1** provide further details and hyper-parameters of the G network. In GAN training, the role of the D network is to drive G to generate data distributed like in the training set. In idpGAN, D receives as input an example ***x*** and an amino acid sequence ***a*** and returns a scalar value corresponding to the probability of the combination being real (that is, from the training set). The input ***x*** represents a protein conformation. It is a vector containing the values of the upper triangle of the interatomic distance matrix calculated from coordinates ***r***. Since interatomic distances are E(3) invariant with respect to transformations of atomic coordinates, using ***x*** as input to D makes idpGAN training invariant to translations, rotations and reflections of the input conformations, an important requirement in 3D molecular generative models ^25^. As the CG representation that we use is not chiral, we can allow reflection invariance in our model. To train idpGAN, the D network must process inputs of variable sizes, that is, data from proteins with variable lengths. Although we experimented with different network architectures with this ability ^32^, we could not identify a solution resulting in stable GAN training. Instead, we found that simple multilayer perceptrons (MLPs) gave good results. Since MLPs take fixed-size input, we employed four MLPs discriminators (each accepting data from proteins with a certain length) along with a scheme for randomly cropping conformations (so that all training proteins could be accepted by one of the MLPs). The idea of using one G and multiple D networks in GANs has been explored elsewhere ^33,34^ and we found it to work well in practice. The “Methods” section provides details of this strategy.

IdpGAN was trained on conformations from MD simulations of a set of 1966 IDPs. These IDPs were obtained from the DisProt ^35^ database and have lengths ranging from 20 to 200 residues (cf. “Methods”). We note that some of the IDPs are actually intrinsically disordered regions (IDRs) in larger proteins, which is not considered here. The MD simulations were performed using a residue-level (Cα-based) CG model that was recently developed in our group (cf. “Methods”). When building the training set, our aim was to obtain sufficient data to span a significant portion of the IDP sequence space, so that our model could learn general rules relating sequences and conformational variability that can be transferred to new sequences.

### IdpGAN evaluation on a test set of IDPs

Since idpGAN is a conditional generative model, once it has been trained, it can generate conformations for proteins of arbitrary sequences. To evaluate whether, for proteins unseen in the training set, the conformational ensembles generated by idpGAN recapitulate the ones observable in MD simulations, we evaluated our model on CG MD data for a set of 31 selected IDPs, named *IDP_test*, that have no similar sequences in the training set (cf. “Methods”). We also compared against data from CG MD simulations for poly-alanine (polyAla) chains with lengths from 20 to 200 as a random linear polymer model without sequence-specific interactions to study whether idpGAN could provide better approximations. To evaluate idpGAN on a test protein, we generated 10,000 conformations. The same number of conformations was randomly extracted from the corresponding MD data and from the polyAla ensembles.

Examples of generated ensembles for three selected *IDP_test* proteins (*his5, protac* and *htau23k17*) are shown in **Fig. 2** (the remaining proteins are shown in **Supplementary Figs. 2** to **7**). Sample illustrations of generated 3D conformations along with their nearest neighbor in the MD data demonstrate that the conformations appear qualitatively “realistic”. Residue contact maps are also matched closely in the examples. In the CG model that we study, specific amino acid sequences influence protein dynamics and give origin to contact maps with patches of relatively lower or higher contact probabilities. The goal of idpGAN was to capture these sequence-specific patterns. In the case of *protac*, there are low-probability regions in the contact map caused by stretches of positively charged amino acids repelling each other (**Supplementary Table 1**). The map generated by idpGAN reproduces a very similar distinctive pattern, even though *protac* (or a similar IDP) was not present in the training data set. This clearly illustrates that idpGAN learned transferable residue-specific interaction patterns from the training MD data. Finally, **Fig. 2** also shows radius-of-gyration and energy distributions (based on the CG model energy function) from the idpGAN-generated models in good agreement with the MD-generated ensembles. This indicates that the models are of practical value in estimating radius-of-gyration distributions and that they are of high structural quality without clashes or other significant violations of stereochemical constraints.

**Table 1:** Evaluation of idpGAN for *IDP_test* set.

| Method | *MSE_c* | *MSE_d*<br>[nm<sup>2</sup>] | *aKLD_d* | *EMD-dRMSD*<br>[nm] | *KLD_r* | *MED*<br>[kcal/mol] |
| --- | --- | --- | --- | --- | --- | --- |
| polyAla | 1.10<br>± 0.34 | 0.55<br>± 0.10 | 0.14<br>± 0.02 | 0.73<br>± 0.05 | 0.53<br>± 0.09 | - |
| idpGAN | 0.18<br>± 0.08 | 0.02<br>± 0.004 | 0.01<br>± 0.001 | 0.56<br>± 0.04 | 0.02<br>± 0.002 | 5.58<br>± 2.48 |
| null-MD | 0.09<br>± 0.06 | 0.0003<br>± 0.00004 | 0.006<br>± 0.001 | 0.543<br>± 0.034 | 0.006<br>± 0.001 | -0.08<br>± 0.10 |
| idpGAN<br>no-stereo<sup>a</sup> | 0.22<br>± 0.09 | 0.01<br>± 0.003 | 0.01<br>± 0.001 | 0.55<br>± 0.04 | 0.01<br>± 0.001 | 67410<br>± 34340 |

Average values are reported along with standard errors for all the proteins in the set ( $N = 31$ ).
<sup>a</sup>idpGAN trained without the stereochemical term in the generator loss.

**Fig. 2.**
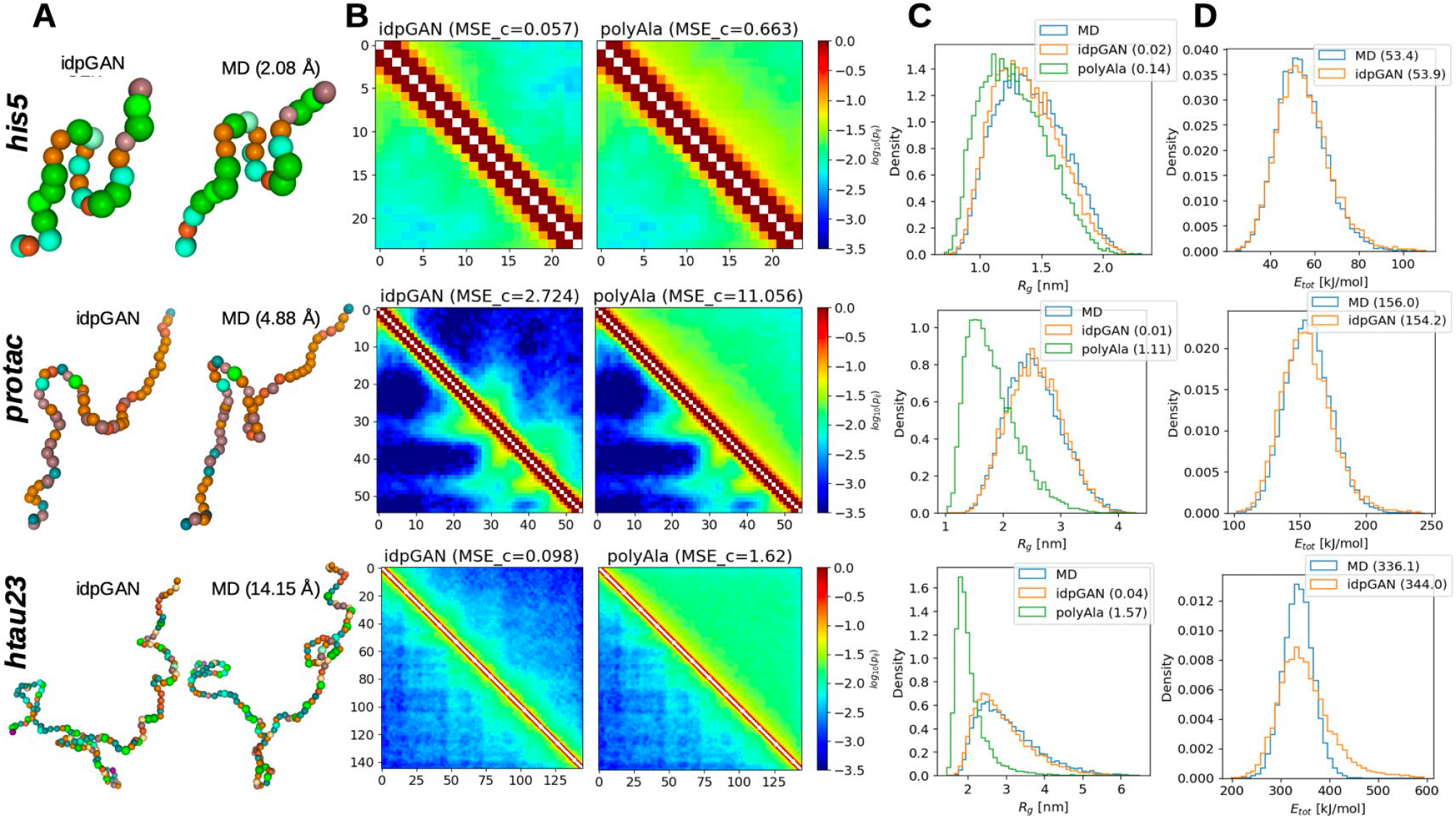
Examples from idpGAN for three IDP_test proteins. Each row of the image shows data for *his5* (*L*=24), *protac* (*L*=55), and *htau23k17* (*L*=145), respectively For each protein, the following information is reported: **A** sample conformation from idpGAN (left) and the nearest neighbor in terms of dRMSD (cf. “Methods”) in MD data (right). The dRMSD value is shown in brackets. The 3D conformations were rendered with Nglview ^36^. **B** idpGAN (left) and polyAla (right) contact maps are shown in the upper triangles of the images. The MD-generated maps are shown in the lower triangles. The values shown along the horizontal and vertical axes are residue indices. The *MSE_c* scores with the MD maps are shown in brackets. **C** Radius-of-gyration distributions for the MD (blue), idpGAN (orange) and polyAla (green) ensembles. *KLD_r* values are shown in brackets. **D** Total potential energy distributions for the MD and idpGAN ensembles based on the CG energy function. The median values of the data are shown in brackets.

For a quantitative evaluation of idpGAN’s performance, we turned to a number of metrics that we calculated across all *IDP_test* proteins (**Fig. 3** and **Table 1**). The same metrics were also calculated for the corresponding polyAla ensemble to provide a random polymer baseline. Another baseline was calculated by drawing 10,000 snapshots from an additional independent long MD run (cf. “Methods”) and comparing those to the MD snapshots from the same reference simulations used for evaluating idpGAN. This baseline essentially captures the variability of sampling between different MD simulation trajectories of the same system since the simulations are of finite length and sampling is likely incomplete. For all these metrics (described in detail in “Methods”), scores closer to zero reflect better approximations of the MD reference ensembles.

**Fig. 3.**
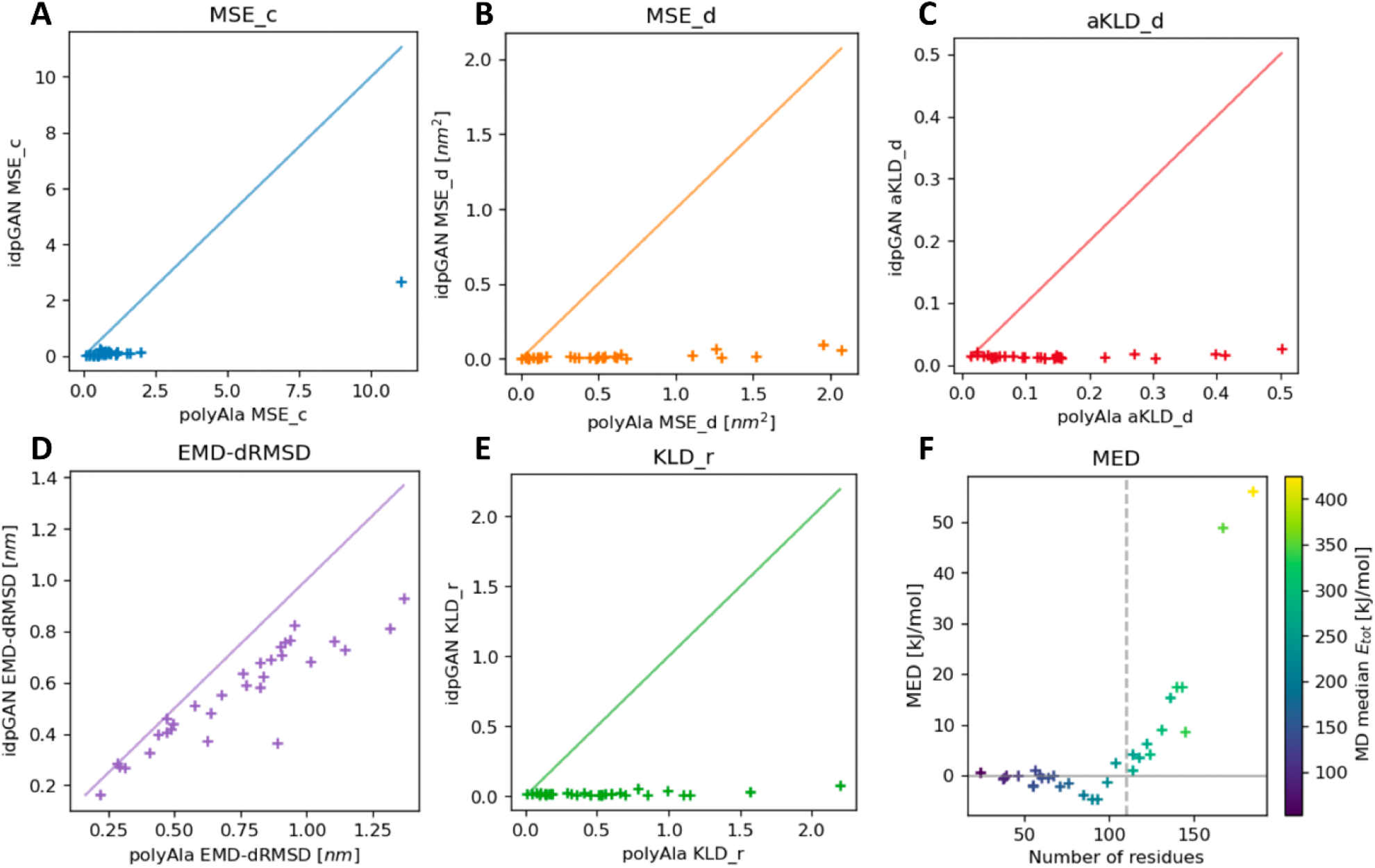
Evaluation of idpGAN and polyAla ensembles for approximating MD data. Results are reported for *IDP_test* set protein (*N* = 31). **A, B, C, D**, and **E** show the values of *MSE_c, MSE_d, aKLD_d, EMD-dRMSD* and *KLD_r*, respectively, obtained by polyAla (x-axis) and idpGAN (y-axis) for all the proteins in the set. Lower values indicate a better performance in approximating MD ensembles. *MED* values as a function of protein length are shown in **F** with markers colored according to the median potential energy of proteins in the MD ensembles. The dashed vertical line represents the maximum crop length used in idpGAN training (*L* = 110).

The first metric we considered is the *contact mean squared error* (*MSE_c*) which quantifies differences in residue contact maps. *MSE_c* values from idpGAN ensembles are close to zero for almost all proteins in the *IDP_test* set, with the example *protac* discussed above (*MSE_c* = 2.89) actually being the largest outlier. In contrast, polyAla ensembles have much larger *MSE_c* values. As the next metric we analyzed average distances in the generated distance matrices according to the *distance mean squared error* (*MSE_d*). Again, the idpGAN-generated average distance maps closely resemble the MD ones (see **Supplementary Fig. 8**), and their *MSE_d* scores are much better than those obtained from polyAla ensembles.

We then evaluated further how well idpGAN models not just distance averages but distance distributions based on the *average Kullback-Leibler divergence for distance distributions* (*aKLD_d*). Again, there was close agreement between idpGAN and MD ensembles, and much better distributions can be obtained with idpGAN than from polyAla samples. This is further illustrated by randomly selected histograms for interatomic distance data (see **Supplementary Fig. 9**).

We continued to test whether idpGAN correctly captured not just pairwise distributions but correlations between multiple pairs. To that extent, we compared multi-dimensional joint distributions comprising all interatomic distances in proteins. To approximate their divergence between different ensembles we used the *earth mover’s distance via distance root mean square deviation* (*EMD-dRMSD*) metric, based upon the approximation in ^37^. As **Fig. 3** shows, there is some divergence between idpGAN and MD distributions in this rather stringent metric, but the average EMD-dRMSD value of idpGAN for the IDP_test set (0.556 nm) is again lower as what is obtained with polyAla data (0.733 nm). Moreover, there is a similar degree of divergence when snapshots are taken from a separate long MD trajectory and compared with the reference MD ensemble (**Supplementary Fig. 10**). This suggests that the larger divergence in this metric is more likely due to incomplete sampling in the reference MD ensemble than due to poor performance of idpGAN.

Finally, energies were compared in terms of *median energy differences* (*MED*). While the distributions do not match perfectly, there is considerable overlap between them and *MED* values are small, on average 5.6 kJ/mol for the proteins in the *IDP_test* set. The differences increase in value as proteins become larger and as the average values of the energies themselves increase. We note that it was crucial to include a stereochemical term in the idpGAN generator loss function (cf. “Methods”) to obtain such conformations with low energies. We also trained and evaluated an idpGAN version in which this term was not used in the G network training objective. According to most evaluation metrics, the performance of this ablated model does not change much (**Table 1**), but the average MED values are much larger as a result of clashes between residue beads, since the short-range energy term in the FF is very sensitive to close contacts (**Supplementary Fig. 11**). Therefore, it seemed necessary to train our GAN model by including physical constraints to capture fine stereochemical aspects.

Taken together, these results show that idpGAN generates energetically-stable conformations and captures the variability of the ensembles in MD data and their amino acid sequence-specific characteristics in a transferable manner.

### Evaluation on a large set of IDPs

We also evaluated idpGAN on the *HB_val* set, a larger set of IDPs obtained through a form of cross-validation in our training set (cf. “Methods”). The evaluation confirms the same trends observed for the *IDP_test* set (**Fig. 4** and **Table 2**), that is, idpGAN provides very good approximations of the MD reference data and much better approximations with respect to random polymer ensembles from polyAla according to all of the metrics.

**Table 2:** Evaluation of idpGAN for *HB_val* set.

| **Method** | ***MSE_c*** | ***MSE_d***<br>**[nm<sup>2</sup>]** | ***aKLD_d*** | ***EMD-dRMSD***<br>**[nm]** | ***KLD_r*** | ***MED***<br>**[kcal/mol]** |
| --- | --- | --- | --- | --- | --- | --- |
| polyAla | 0.81<br>± 0.05 | 0.32<br>± 0.03 | 0.10<br>± 0.005 | 0.35<br>± 0.02 | 0.32<br>± 0.02 | - |
| idpGAN | 0.14<br>± 0.02 | 0.02<br>± 0.002 | 0.02<br>± 0.001 | 0.37<br>± 0.01 | 0.02<br>± 0.001 | 0.83<br>± 0.50 |
| idpGAN<br>no-stereo<sup>a</sup> | 0.16<br>± 0.02 | 0.01<br>± 0.002 | 0.01<br>± 0.001 | 0.37<br>± 0.01 | 0.02<br>± 0.001 | 84371<br>± 15744 |

Average values are reported along with standard errors for all the proteins in the set ( $N = 1021$ ).
<sup>a</sup>idpGAN trained without the stereochemical term in the generator loss.

**Fig. 4.**
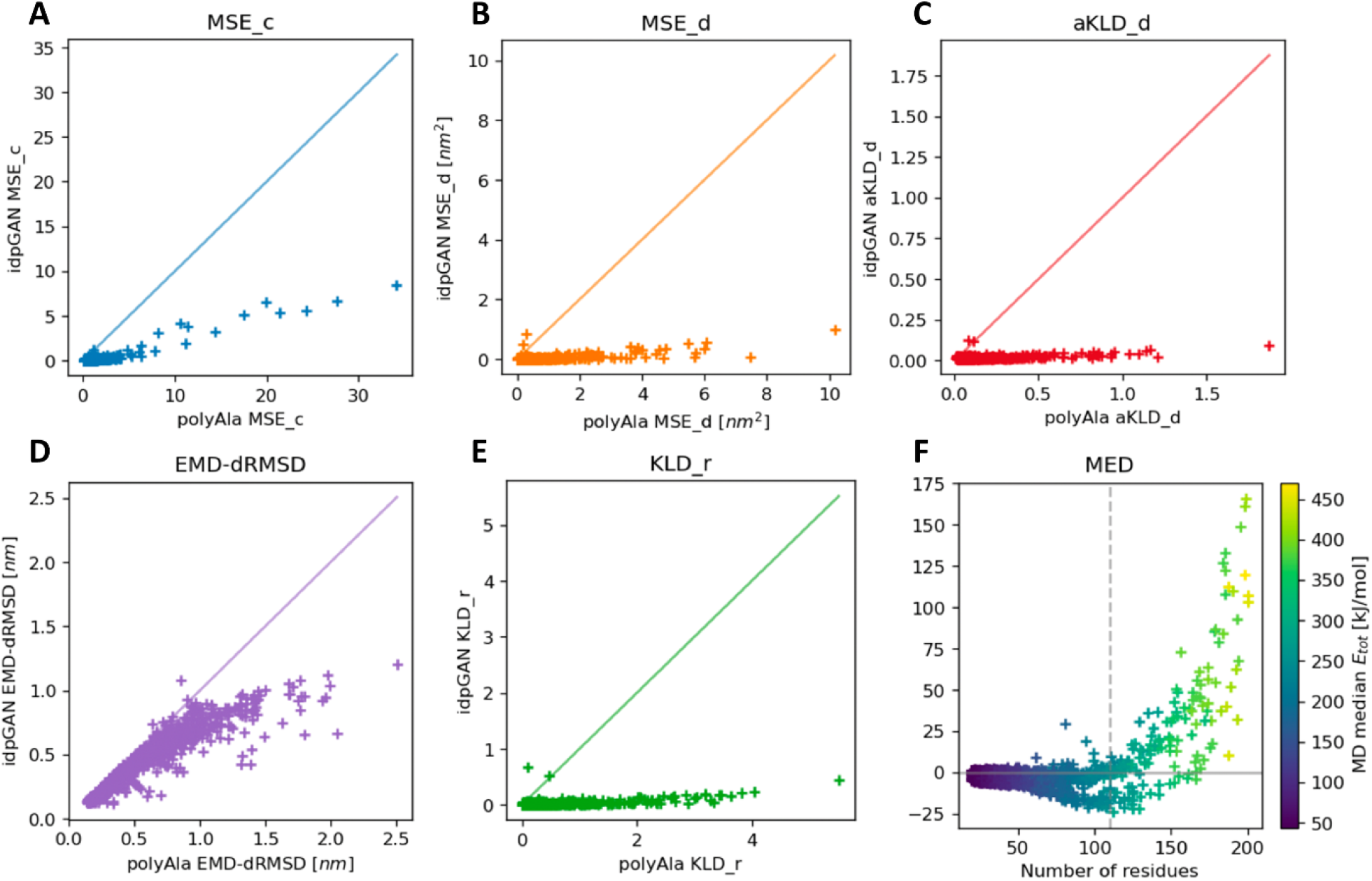
Evaluation of idpGAN and polyAla ensembles for approximating MD data. Results are reported for *HB_val* set protein (*N* = 1021). Metrics are shown as in **Fig. 3**.

Evaluating idpGAN on more IDPs permitted us to examine how its performance is affected by specific sequence features. Protein length appears to have an important impact. **Fig 4F** shows that *MED* values of IDPs with lengths above 150 frequently surpass 50 kcal/mol and the values increase for longer IDPs. Note that the maximum IDP crop length used in idpGAN training is 110. Therefore, it seems that the model has difficulties in generating energetically stable conformations for proteins with lengths above the maximum one used in training. In **Supplementary Fig. 12D**, we show the distribution of each energy term of *DP02478r001*, the *HB_val* IDP with the highest *MED* value (166.1 kcal/mol). The figure shows that, in the generated ensembles, the terms with higher median values with respect to the reference distributions are the bond length, bond angle and short range interaction ones. Despite high *MED* values, the distributions of the corresponding geometrical features (**Supplementary Fig. 12E**) and several properties of the ensemble (such as the contact map and the distribution of radius-of-gyration, see **Supplementary Fig. 12A** to **C**) appear to be well-captured by the model. The high energies are explained by the fact that in our FF (like in numerous molecular mechanics FFs) these three energy terms are quadratic or higher polynomial functions of their input geometrical features. For this reason, even small divergences in the distributions of the features lead to large energies. Other evaluation metrics are also influenced by protein length, although for most of them the effect is weaker (**Supplementary Fig. 13**).

This apparent limitation of idpGAN when applied to longer proteins could be overcome by training with longer crops or by adding a neural-network based refinement post-processing step as in ^24^. Even though we did not explore these possibilities due to time and computational constraints, we believe that they could be valid strategies to further improve idpGAN performance.

### IdpGAN sampling speed

One of the main goals of idpGAN is to obtain better computational efficiency than MD. Sampling with a GAN is very fast since it only takes a forward pass of the G network, which can be performed in a highly efficient way with modern deep learning libraries. For proteins with lengths below 150 residues, it typically takes less than 1 s to generate ensembles containing a number of independent conformations in the order of thousands (**Supplementary Fig. 14**).

To compare the sampling speed of MD simulations and the idpGAN generator, we measured the GPU time used by both to generate enough samples to recover the distribution of the radius-of-gyration observed in 5 μs MD runs, which we note are already highly efficient as they involve a CG model run on a GPU. In **Fig. 5A**, we plot the *KLD_r* of ensembles from the G network (orange data points) and MD ensembles (blue) with increasing numbers of samples when they are compared with the long MD run ensembles. For both the G network and MD simulations, *KLD_r* improves as the methods sample more conformations (by consuming more GPU time) and ultimately tends to zero for MD. For *his5, protac* and *htau23k17*, the computational time it takes for the G network to reach a plateau in *KLD_r* (referred to as *t*_*gen*_) is always less than 3 s. The time it takes for an MD simulation to reach the same *KLD_r* values (referred to as *t*_*MD*_) is always above 250 s. While idpGAN may not perfectly recover the long MD run distributions, it can provide close approximations as evaluated by *KLD_r* (**Fig. 2C**) and accomplish this orders of magnitude faster than what MD is able to achieve. Similar trends are confirmed for the rest of the *IDP_test* proteins (see **Fig. 5B**). The efficiency of idpGAN relative to MD simulations is better for shorter proteins since the G network can generate more conformations in parallel on a GPU.

**Fig. 5.**
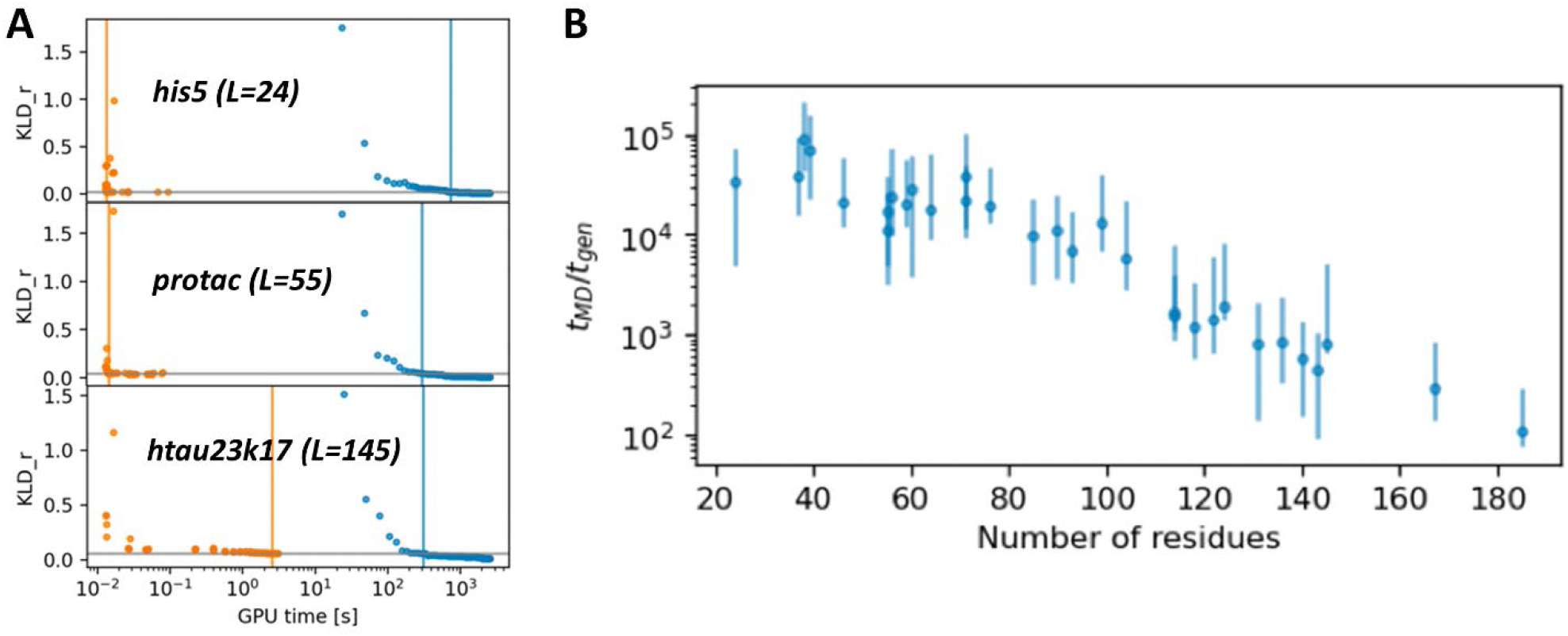
Evaluation of idpGAN sampling efficiency compared to MD. **A** *KLD_r* between the idpGAN and reference MD ensembles (orange data points) and between short MD and reference MD ensembles (blue data points) as a function of GPU time. Orange vertical lines indicate the GPU time (*t*_*gen*_) at which the generator reaches a plateau *KLD*_*r*_*top*_ value, which is marked by a gray horizontal line. The blue vertical lines indicate the GPU time (*t*_*MD*_) needed by MD to surpass the *KLD*_*r*_*top*_ value. Data is shown for *his5, protac*, and *htau23k17* proteins with t_gen_ values of 1.4 × 10^−2^ s, 1.5 × 10^−2^ s, and 2.6 s and *t*_*MD*_ values of 656.1 s, 269.3 s, and 309.6 s, respectively, for the three proteins. **B** *t*_*MD*_ /*t*_*gen*_ ratio for all proteins of the *IDP_test* set are plotted as a function of protein length. The values are the mean of 10 runs, and the bars report minimum and maximum values across the runs.

### Modeling the Cα trace of all-atom trajectories

The idpGAN model described above was trained with CG protein conformations, or more precisely with conformations generated with simulations of a CG model, which may be of more limited value when considering biological applications. To establish if idpGAN can be adapted to model all-atom protein conformational ensembles, we re-trained the model with data from all-atom MD simulations. There are three aspects when considering all-atom trajectories: 1) To capture atomistic details, a network with much greater capacity would be needed. As we are focusing here on a proof-of-principle, we avoid this challenge by considering only the Cα atoms extracted from all-atom trajectories; 2) Because all-atom simulations are very expensive, especially for highly dynamic and extended IDPs, we focus on only one system, α-synuclein (140 residues), for which we have simulation trajectories from previous work ^30^. Therefore, the re-trained model is an unconditional generative model that is not transferable to other sequences; 3) The high-resolution interaction potential used in all-atom simulations gives rise to more complex features of the IDP ensemble (**Fig. 6**) and the main test here is in fact whether the idpGAN architecture can capture such detailed features faithfully.

**Fig. 6.**
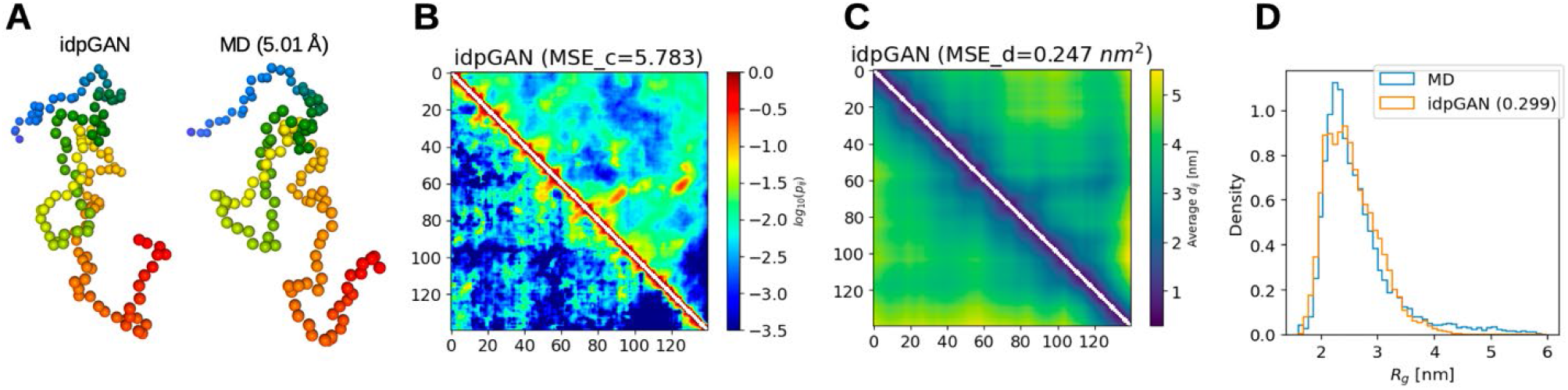
Modeling of the conformational ensemble of α-synuclein from all-atom simulations using idpGAN. **A** 3D structures of a generated conformation (left) and the nearest neighbor in terms of dRMSD in MD data (right). **B** idpGAN contact map (in the upper triangle of the image) confronted with the MD map (lower triangle). Their *MSE_c* score is shown in brackets. **C** idpGAN average distance map (upper triangle) confronted with the corresponding MD map (lower triangle). Their *MSE_d* score is shown in brackets. **D** Radius-of-gyration distributions for the MD (blue) and idpGAN (orange) ensembles with the *KLD_r* value shown in brackets.

It turned out that our original idpGAN network was not able to model this more complex data, as training was highly unstable. To successfully adapt our model, we had to increase its capacity and change its training objective to stabilize learning (cf. “Methods”). **Fig. 6** shows validation of a model that was trained based on snapshots from one 2 μs all-atom simulation by comparing against MD ensembles extracted from two different, independent trajectories ^30^. The generated conformations appear as realistic and the contact maps show the same overall structure, but with moderate deviations from the MD-based contacts (*MSE_c* = 5.78) and some differences in the detailed features that indicate some overfitting to features specific to the training trajectory, such as contacts between the regions around residues 60 and 130 (**Supplementary Fig. 15**). The average distance maps are also similar (*MSE_d = 0*.*25* nm^2^) and the generated distance distributions are overall correct (**Supplementary Fig. 16**). Finally, the radius-of-gyration distributions also share similar forms (*KLD_r* = 0.30).

These results show that a network like idpGAN has the potential to model all-atom protein conformational ensembles. With additional modifications in the neural network and training process together with using larger and more diverse training data sets, an idpGAN-like system may be extended to accurately model dynamics at the all-atom level for arbitrary systems.

## DISCUSSION

The machine learning model idpGAN is presented here to demonstrate that one can generate realistic conformational ensembles of protein structures in a highly efficient manner. Most importantly, the GAN-based model can generate conformations for previously unseen proteins, with chemically reasonable stereochemistry and with favorable and correctly distributed energetics. The model directly generates structures that can make up a complete ensemble of energetically favorable conformations. There is no physics-based iterative sampling, which makes the approach extremely fast, but a drawback is the loss of any dynamic information, since each generated conformation is completely independent from any others. However, knowing the conformational ensemble for a given system would allow dynamics and kinetics to be recovered via simulation-based re-sampling methods ^38^.

Training of the model relied on conformational sampling extracted from MD simulations. The advantage is that such training data can be generated relatively easily if it is not yet available. As a consequence, the machine learning model mirrors the physical realism as well as any artefacts the simulations may provide. Ideally, one would like to learn from experimental data, but unfortunately there is not much high-resolution data on conformational sampling, especially for more flexible elements in large macromolecules, such as proteins. One strategy in the future may be to train using simulations with multiple force fields and introduce additional constraints to incorporate experimental knowledge during the training to increase the physical realism. Another issue is that the MD simulations used for training need to be long enough to recover the underlying equilibrium distributions of conformations, as training with incomplete data could result in a generative model learning the wrong distribution. This is a difficult-to-meet requirement, although a solution for this problem might come in the form of Boltzmann generators ^27^.

An important point of any machine learning approach is transferability without which there is limited utility. We demonstrate that our conditional GAN model learned sufficiently general features to be able to predict correct ensembles for sequences not included in the training data. The current model works well for sequences up to and perhaps slightly beyond the longest sequence in the training set, but there is some deterioration in structure generation for much longer sequences as shown by the fact that idpGAN does not capture the correct energy distributions for longer proteins. Presumably this could be overcome by expanding the training set to include larger proteins.

In our GAN framework, the use of multiple simple neural networks as discriminators is probably inefficient and likely limits performance. We believe that employing models with stronger inductive biases as discriminators, such as graph neural networks ^32^, will be one of the keys to improve our method. Additionally, our training objective uses a very simple physical-based term to improve the stereochemical quality of the generated conformations. Using more sophisticated and effective ways of including prior physical-based constraints in the learning process of the model will also be important for improving its performance. From a generative model point of view, although GANs are powerful models, their training instability ^39^ creates practical challenges. Therefore, other kinds of generative models may improve our approach. Research in generative modeling is certainly flourishing ^40,41^ and new methods, such as probabilistic diffusion models, are continuously being developed as possible alternatives.

For practical reasons, *i*.*e*. ease of generating simulation data and ease of training, our model was principally destined for CG conformations. However, we show that the approach can be extended to all-atom representations. To limit computational complexity during training and because MD training data is scarce, we focused here on modeling the Cα traces of all-atom simulations of only one IDP system, α-synuclein, for which extensive MD data was already available. As the number of atoms to model increases, the training time of a generative model becomes much longer. Training a model on large-scale all-atom MD protein datasets would require significant resources on its own. Since, at the present time, machine learning method development is largely empirical, requiring numerous trial and error iterations, the computational burden may limit the use of this type of approach on all-atom protein data in the near future. However, there is no fundamental reason that our approach could not be extended to develop a broader model that eventually predicts conformational ensembles of any protein at the atomistic level, given that algorithmic and hardware capabilities advance accordingly. Nevertheless, depending on the application, the current approach focusing on a CG representation may already be sufficient, for example to generate approximate radius-of-gyration distributions of IDPs for interpretation of experimental data ^30^.

We demonstrate that once a generative model is trained on MD data, it can sample from the underlying distributions orders of magnitude faster, and even much faster than CG methods that are traditionally used to accelerate conformational sampling. However, a central advantage of physics-based approaches, like MD, is that changes in physical conditions (temperature, pH) or chemical composition (different solvents or the presence of other solutes) are, at least in principle, easily incorporated. A machine learning model not trained with data reflecting such external variations, will not be able to provide any insights on such factors. On the other hand, generating comprehensive training data for a variety of conditions and a variety of systems is probably not practical at the current time. A model based on neural networks incorporating stronger inductive biases for molecular data ^32^ could mitigate the dependence on the amount of training data. However, either learning all of the physics or re-introducing it as part of the machine learning model will be the most significant challenge in achieving a universally applicable conformational ensemble generator.

## METHODS

### Training and test sets

The training set of idpGAN consists of CG MD data for a series of IDPs. We started by defining a test set of 31 IDPs, named *IDP_test* (**Supplementary Table 1**), by selecting proteins based on availability of their experimental radius-of-gyrations and with a goal of covering distinct sequence lengths. We note that some of these proteins are actual IDPs under biological conditions while others are natively folded proteins that were characterized in the presence of denaturant. The training set was then constructed by selecting all IDPs from DisProt ^35^ (version *2021_06*), a database of protein disordered regions, with lengths ranging from 20 to 200 ns. We note that many IDPs are actually intrinsically disordered regions in larger proteins, which we neglect here. To ensure that peptides with similar sequences are not present in both training and test sets, we then removed 32 IDPs from the initial training set because of sequence similarity with proteins in *IDP_test*. Here, we define a query sequence as “similar” to a training sequence if they have an E-value < 0.001 in a phmmer search ^42^ scanning the training set with default parameters. This yielded a final training set with 1966 IDPs (see **Supplementary Fig. 17** for the distribution of their lengths and amino acid composition).

### Coarse-grained molecular dynamics simulations

To obtain conformational data for training and testing idpGAN, we ran MD simulations for all the training and test set IDPs using a recently developed CG model from our group. In this model, each residue is represented as a single spherical particle located at the Cα atom of a given residue. The potential form is given by:

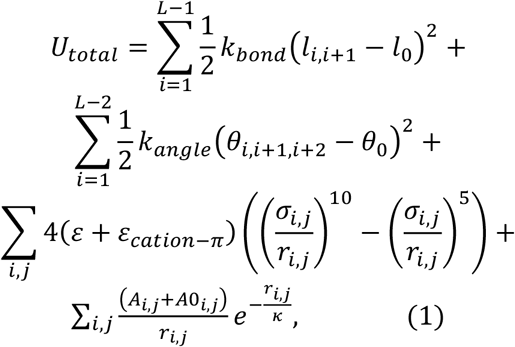

where *L* is the number of residues in a protein. The bonded parameters are as follows: *l*_*i,i*+1_ is the distance between two neighboring residues, with the spring constant *k*_*bond*_ = 4,184 kJ/(mol·nm^2^), *l*_0_ = 0.38 nm is the equilibrium bond length; θ_*i,i*+1,*i*+2_ is the angle between two subsequent Cα beads, with an angle spring constant of *k* _*angle*_ = 4.184 kJ/(mol·rad^2^), and an equilibrium angle θ_0_ = 180°. The remaining terms refer to non-bonded interactions: *r*_*i,j*_ is the inter-residue distance for residues not connected via bonds, *σ*_*i,j*_ = *σ*_*i*_ + *σ*_*j*_ where *σ*_*i*_ was determined as the radius of a sphere with an equivalent volume of a given residue, ε is set to 0.40 and 0.41 kJ/mol for polar and non-polar residues, respectively, ε_cation−π_ is set to 0.3 kJ/mol to augment interactions between basic residues (Arg/Lys) and aromatic residues (Phe/Tyr/Trp); A_*i,j*_ = A_*i*_ × A_*j*_ describes long-range interactions with 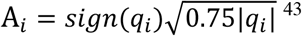 using charges q_*i*_ = +1 for Arg/Lys, q_*i*_ = −1 for Asp/Glu, and q_*i*_ = 0 for all other residues; A0_*i,j*_ = A0_*i*_ × A0_*j*_ describes the repulsion between polar residues due to solvation with A0_*i*_ being 0.05 for polar and 0 for non-polar residues, respectively.

For all of the IDPs as well as the polyAla reference, we ran simulations with OpenMM 7.7.0 ^44^ using the CG interaction potential described above. Langevin dynamics was used with a friction coefficient equal to 0.01 ps^−1^. A short equilibration was performed initially with 5,000 steepest descent minimization steps followed by 20,000 steps of molecular dynamics with a 0.01 ps time step. For production runs we increased the time step to 0.02 ps. Non-bonded interactions were calculated considering periodic boundary conditions and interactions were truncated at 3 nm. Bonded residues were excluded in non-bonded interaction evaluations. Individual protein chains were simulated in a cubic box of side 300 nm at 298 K. For all proteins we ran five separate trajectories over 1,000 ns and one additional longer trajectory over 5,000 ns for the proteins in the *IDP_test* set. Coordinates were saved every 200 ps giving a total of 5 × 5,000 = 25,000 and 1 × 25,000 = 25,000 trajectory snapshots for each protein. Initial random coordinates for each chain were obtained using a custom Python script. The topology was then generated using the MMTSB Tool Set ^45^ and CHARMM v44b2^46^.

### IdpGAN training objective

The learning process of a GAN involves the training of two neural networks, G and D ^29,47^. IdpGAN is a conditional GAN ^48^, trained with the non-saturated GAN objective ^29,49^. The discriminator loss, denoted as *L*_*D*_, is:

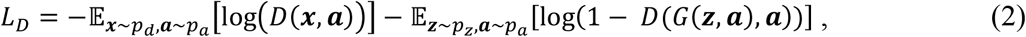

where *p*_*b*_ is the distribution in the training set of examples ***x*** (that describe molecular conformations), *p*_*a*_ is the distribution of examples ***a*** (that represent amino acid sequences), and *p*_*z*_ is the prior from which *z* values are sampled (see below). The discriminator output *D*(*x*, ***a***) is a scalar from 0 to 1 and represents the probability of a sample ***x*** with sequence ***a*** to be real. The output of *G*(*z*, ***a***) is a generated conformation ***x*** for a protein with sequence ***a***.

For the generator loss, we modified the original non-saturated GAN loss by including a term *E*_*C*_, inspired by a term in the AF2 objective ^10^, to reduce the number of steric clashes between “non-bonded” atoms, which we define as atoms in residues with a difference in position of 3 or more. The term takes as input a vector ***x*** (which stores all interatomic distances in a conformation, see below) and is expressed as:

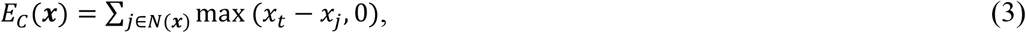

where *N*(*x*) is the set of indices for all “non-bonded” distances in ***x*** and *x*_*t*_ = 0.59 nm is a threshold corresponding to the 0.1 percentile of the training set “non-bonded” distances. The generator loss, denoted as 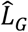, is therefore:

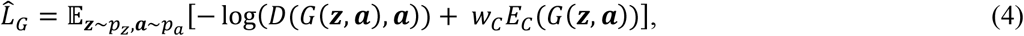

where *w*_*C*_ is a weight that we empirically set to 0.3.

### Generator and discriminator networks

We implemented idpGAN neural networks using the PyTorch framework ^50^. The G network has a transformer architecture ^31^. For a protein of length *L*, the network takes as input: (1) a tensor 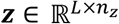 whose values are randomly sampled from a gaussian *N*(0, I); (2) a tensor ***a*** ∈ ℝ^*L*×20^ storing one hot encodings for the amino acid types of the protein. The output of the network is a tensor ***r*** ∈ ℝ^*L*×3^, representing the 3D coordinates of the protein Cα atoms. Please refer to **Supplementary Fig. 1** for a detailed description of the G network.

As discussed in the “Results” section, we employ 4 MLPs discriminators. Each discriminator has the same architecture and hyper-parameters, with a non-trainable standardization layer, three linear layers with spectral normalization ^39^ to regularize GAN training, two leaky ReLU non-linearities and a final sigmoid activation (see **Supplementary Table 2**). The input of a discriminator is composed as follows: starting from a conformation ***r***, we compute its distance matrix ***X*** ∈ ℝ^*L*×*L*^, extract its upper triangle (excluding its zero-filled diagonal) and flatten it to a vector ***x*** ∈ ℝ^*L*(*L*−1)/2^. We then process ***x*** through a standardization layer where each value *x*_*i*_ (representing a distance between two Cα atoms with sequence separation *k*) is converted via 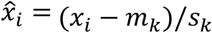, where *m*_*k*_ and *s*_*K*_ are the mean and standard deviation in the training set for distances between Cα atoms with separation *k*. To provide amino acid sequence information, 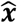 is concatenated to a flattened version of a (a 20*L*-dimensional vector). The resulting *L(L-1)/2+20L*-dimensional vector is the input of the first linear layer of the MLP.

### Training process

To train idpGAN, we chose a set of 4 length values *L*_*train*_ = (20, 50, 80, 110). For each value, we employed a MLP discriminator that only takes as input proteins crops with that length. To make use of all IDPs, at the beginning of each training epoch we adopt the following strategy:

1. First, each IDP with a length value present in *L*_*train*_ is associated with the corresponding MLP.
2. For IDPs with length values not in *L*_*train*_, we crop them to the closest value in *L*_*train*_. In this way, the following number of IDPs are assigned for different *L*_*train*_ values: 1,070 for 20, 317 for 50, 246 for 80, and 279 for 110.
3. Because of the distribution of training IDP lengths (see **Supplementary Fig. 17**), lower *L*_*train*_ values have more counts than the rest. To avoid unbalanced training, we associate *c*_*max*_ = 1,070 IDPs (the highest count obtained in the previous step) to all *L*_*train*_ values. To do that, we randomly sample all training IDPs and assign each to a random *L*_*train*_ value, until all MLPs are associated with *c*_*max*_ IDPs.
4. Each time an IDP is assigned to a MLP, we randomly sample *n*_*frames*_ = 1,750 frames from its MD data. When an IDP is cropped, each frame is randomly cropped selecting different starting and ending residues. This random selection scheme at each epoch effectively works as a form of data-augmentation.

By using this strategy, we have |*L*_*train*_ | × *c*_*max*_ × *n*_*frames*_= 4 × 1,070 × 1,750 = 7,490,00 training MD frames per epoch.

To optimize the idpGAN objective, we use Adam optimizers ^51^ (with *β*_*1*_ = 0.0 and *β*_*2*_ = 0.9 hyper-parameters) and employ learning rates of 0.00025 and 0.0004 for G and all D networks (each D network has its own optimizer).

For training, we use a batch size of 196, with batches containing crops having the same number of residues. The training of an idpGAN model lasts for 50 epochs, which takes roughly three days on a NVIDIA RTX 2080 Ti GPU. For all runs, we repeat training for 10 times and select the model with the best performance on validation data.

### Evaluation strategy

For initial evaluation of idpGAN, we used the *IDP_test* consisting of 31 IDPs as described above. To evaluate idpGAN on more proteins, we also split the training set into partitions based on sequence similarity ^52^. By performing an all vs. all search with phmmer, we identified 1,021 IDPs that do not have other similar sequences in the set. Removing this subset, which we call the *HB_val* set, the remaining training set would likely be too small to obtain good performance in terms of generalization to arbitrary sequences. Therefore, we randomly split the *HB_val* subset into 5 approximately equally-sized partitions. For each partition, we trained idpGAN with all remaining IDPs, including the four other partitions of the *HB_val* set, and then validated it on the IDPs of the selected partition itself. By repeating this procedure with all partitions, we were able to evaluate the performance of idpGAN on all of the 1,021 proteins of the *HB_val* set.

### Evaluation metrics

To compare idpGAN (or polyAla) ensembles with the reference (MD) ones, we use the metrics described below. In all cases, we generated ensembles with *n*_*eval*_ = 10,000 randomly sampled conformations.

To evaluate contact probabilities, we use the *MSE_c* metric. For a protein of length *L*, it is computed as:

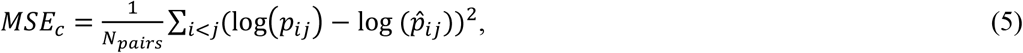

where *N*_*pairs*_ = L(L-1)/2 is the number of residue pairs in the protein, *p*_*ij*_ and 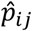 are the contact frequencies for residues *i* and *j* in the reference and generated ensembles respectively (to avoid frequencies equal to zero, we employ a pseudo-count value of 0.01). To define a contact, we use a Cα distance threshold of 8.0 Å.

To evaluate average interatomic distance values, we use the *MSE_d* metric, which is computed as:

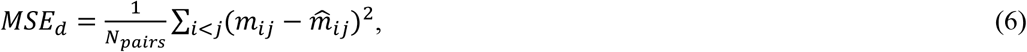

where *m*_*ij*_ and 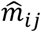 are the average distance values between the Cα atoms of residue pair *i* and *j* in the reference and generated ensembles.

To compare mono-dimensional distributions of several continuous features, we employ an approximation of the Kullback-Leibler divergence (KLD) by discretizing them in the following way. We first take the minimum and maximum value of a feature over the reference and generated ensembles and uniformly split the range in *N*_*bins*_ = 50 bins. We then compute the frequencies of observing values in each bin (with a pseudo-count value of 0.001). Finally, we approximate KLD as:

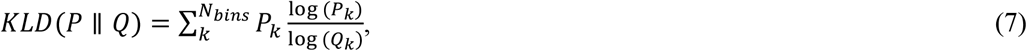

where *k* is the index of a bin and *P*_*k*_ and *Q*_*k*_ are the frequencies associated with bin *k* for the reference and generated ensembles respectively.

To compare pairwise interatomic distance distributions, we use the *aKLD_d* metric, which is expressed as:

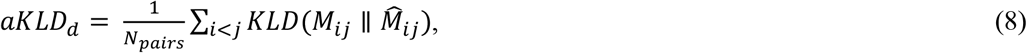

where *M*_*ij*_ and 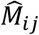are the distance distributions between the Cα atom of residue *i* and *j* in the reference and generated ensembles.

To compare the multi-dimensional joint distributions of all interatomic distances in proteins, we approximate their earth mover’s distance (EMD) by considering the distance root mean square deviation (dRMSD) between conformations, inspired by the approximation in ^37^. Given a generated and reference ensemble, we compute dRMSD values for all pairs of conformations between them. The dRMSD for two conformations is:

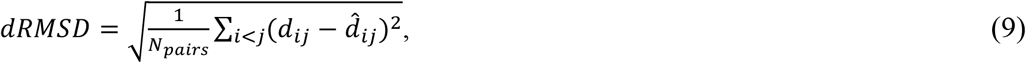

where *d*_*ij*_ and 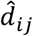 are the distances between Cα of residues *i* and *j* in the reference and generated structures. We then use the Hungarian algorithm to pair each reference conformation to a generated one, to minimize the global dRMSD. The total dRMSD is finally averaged over pairs and reported as the EMD-dRMSD metric.

To compare radius-of-gyration distributions, we use the the *KLD_r* metric expressed as:

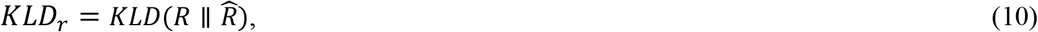

where *d* and *d* are the distributions of radius-of-gyration for the reference and generated ensembles.

### Evaluating sampling efficiency

Sampling efficiency of idpGAN for *IDP_test* proteins was determined as follows:

- We first run a 50,000 ns long MD simulation for a given protein (see above) and we define as *E*_*MD*_ the conformational ensemble containing all 25,000 snapshots in the simulation.
- We then use the G network to sample an increasing number of conformations *n* (from 50 to 15,000). For each generated ensemble *E*_*gen,n*_, we measure its *KLD_r* with *E*_*MD*_. The *KLD_r* values decrease as the number of samples increases, and soon reach a plateau (see **Fig. 5**). If the *KLD_r* does not decrease any more even after generating an additional 1,000 samples, we stop. We denote the minimum *KDL_r* value obtained in this way as *KLD_r*_*top*_ and refer to the time it took for G to generate the corresponding ensemble as *t*_*gen*_.
- Finally, we start extracting an increasing number of snapshots m from the beginning of the long MD simulation (from frame 50 to 25,000). For each MD ensemble *E*_*MD,m*_, we measure its *KLD_r* with *E*_*MD*_. When we find an ensemble that improves over *KLD_r*_*top*_, we denote as *t*_*MD*_ the time taken for the simulation to accumulate the number of samples contained in it.

All runs were performed on NVIDIA RTX 2080 Ti GPUs. When generating samples with idpGAN, we tried to use mini-batches as large as possible to harness the parallel computing capabilities of a GPU. Since the memory requirement of transformers is quadratic with sequence length *L*, we adopted different batch sizes for different *L* values to fit the batches on memory (**Supplementary Table 3**).

### Modeling Cα traces from all-atom simulations

The training data we used for all-atom α-synuclein modeling is a 2 μs simulation containing 10,000 snapshots. For testing we compared with data from two additional independent simulations of the same system. Details of the simulations are given elsewhere ^30^. To evaluate idpGAN, we generated 5,000 samples and employed the evaluation metrics described above.

In this learning task, when using the non-saturated GAN loss (see above), the generator tended to mode-collapse ^47^. By testing other losses, we found that the Wasserstein loss with gradient penalty ^53^ produced stable training. We therefore adopted it with 5 critic iterations and λ = 10. Please refer to ^53^ for details on this loss.

To adapt idpGAN for this more complex data, we also had to modify its G network by increasing its capacity and introducing other small modifications (see **Supplementary Table 4**). α-synuclein has *L* = 140 residues. The input size of the discriminator is constant and therefore we employed a single MLP. Its architecture is similar to the one used for modeling CG conformations (see **Supplementary Table 2**). The discriminator input is a vector of dimension *L*(*L*-1)/2+(*L*-3). The *L*(*L*-1)/2 features account for the distances between all Cα pairs, while the *L*-3 features account for the dihedral angles *θ* between all groups of four consecutive Cα atoms. We included these latter features, because *θ* angles in all-atom protein representations are stereo-specific. By default, idpGAN is reflection invariant, but by including *θ* angles as features to the D network, we can force G to learn the correct mirror images for α-synuclein Cα traces. While training, we adopt a per-feature non-trainable normalization layer, as described for the CG idpGAN version. No conditional amino acid information was inputted since we considered only one system.

We employed the same optimizers described above. We trained idpGAN for 300 epochs using a batch size of 64 and decreased by a factor of 0.5 both G and D learning rates at epochs 100 and 200. We repeated training 10 times and selected the model with the best performance on validation data.

## Data availability

Training set and *IDP_test* sequences and *HB_val* validation splits are available at https://github.com/feiglab/idpgan.

## Code availability

The code for the idpGAN generator, together with its neural network weights are available at https://github.com/feiglab/idpgan. In the repository, we also share Jupyter notebooks with examples on how to use idpGAN to generate and analyze 3D conformational ensembles for user-defined protein sequences.

## ACKNOWLEDGEMENT

This research was supported by National Institutes of Health Grant R35 GM126948.

## AUTHOR CONTRIBUTIONS

GJ, LH, and MF designed the research, GJ performed and analyzed the machine learning work, GVG performed and analyzed the simulations used for training, and all authors jointly discussed the findings and wrote the manuscript.

## REFERENCES

1 Miller, M. D. & Phillips, G. N. Moving beyond static snapshots: Protein dynamics and the Protein Data Bank. J. Biol. Chem. 296, 100749, doi:10.1016/j.jbc.2021.100749 (2021).

2 Otten, R. et al. How directed evolution reshapes the energy landscape in an enzyme to boost catalysis. Science 370, 1442–1446, doi:10.1126/science.abd3623 (2020).

3 Knoverek, C. R., Amarasinghe, G. K. & Bowman, G. R. Advanced Methods for Accessing Protein Shape-Shifting Present New Therapeutic Opportunities. Trends Biochem. Sci. 44, 351–364, doi:10.1016/j.tibs.2018.11.007 (2019).

4 Eisenmesser, E. Z. et al. Intrinsic dynamics of an enzyme underlies catalysis. Nature 438, 117–121, doi:10.1038/nature04105 (2005).

5 Gupta, A. et al. Experimental techniques to study protein dynamics and conformations in: Advances in Protein Molecular and Structural Biology Methods (eds Timir Tripathi & Vikash Kumar Dubey) 181–197 (Academic Press, 2022).

6 Dror, R. O., Dirks, R. M., Grossman, J. P., Xu, H. & Shaw, D. E. Biomolecular simulation: a computational microscope for molecular biology. Annu. Rev. Biophys. 41, 429–452, doi:10.1146/annurev-biophys-042910-155245 (2012).

7 Shaw, D. E. et al. Anton 2: Raising the Bar for Performance and Programmability in a Special-Purpose Molecular Dynamics Supercomputer in: SC ‘14: Proceedings of the International Conference for High Performance Computing, Networking, Storage and Analysis. 41–53 (2014).

8 Abrams, C. & Bussi, G. Enhanced Sampling in Molecular Dynamics Using Metadynamics, Replica-Exchange, and Temperature-Acceleration. Entropy 16, 163–199, doi:10.3390/e16010163 (2014).

9 AlQuraishi, M. Machine learning in protein structure prediction. Curr. Opin. Chem. Biol. 65, 1–8, doi:10.1016/j.cbpa.2021.04.005 (2021).

10 Jumper, J. et al. Highly accurate protein structure prediction with AlphaFold. Nature, doi:10.1038/s41586-021-03819-2 (2021).

11 Baek, M. et al. Accurate prediction of protein structures and interactions using a three-track neural network. Science 373, 871–876, doi:10.1126/science.abj8754 (2021).

12 Pereira, J. et al. High-accuracy protein structure prediction in CASP14. Proteins 89, 1687–1699, doi:10.1002/prot.26171 (2021).

13 Ruff, K. M. & Pappu, R. V. AlphaFold and Implications for Intrinsically Disordered Proteins. J. Mol. Biol. 433, 167208, doi:10.1016/j.jmb.2021.167208 (2021).

14 Thomasen, F. E. & Lindorff-Larsen, K. Conformational ensembles of intrinsically disordered proteins and flexible multidomain proteins. Biochem. Soc. T. 50, 541–554, doi:10.1042/BST20210499 (2022).

15 Oldfield, C. J. & Dunker, A. K. Intrinsically disordered proteins and intrinsically disordered protein regions. Annu. Rev. Biochem. 83, 553–584, doi:10.1146/annurev-biochem-072711-164947 (2014).

16 Mardt, A., Pasquali, L., Wu, H. & Noé, F. VAMPnets for deep learning of molecular kinetics. Nat. Commun. 9, 5, doi:10.1038/s41467-017-02388-1 (2018).

17 Wang, D. et al. Efficient sampling of high-dimensional free energy landscapes using adaptive reinforced dynamics. Nat. Comput. Sci. 2, 20–29, doi:10.1038/s43588-021-00173-1 (2022).

18 Husic, B. E. et al. Coarse graining molecular dynamics with graph neural networks. J. Chem. Phys. 153, 194101, doi:10.1063/5.0026133 (2020).

19 Bond-Taylor, S., Leach, A., Long, Y. & Willcocks, C. G. Deep Generative Modelling: A Comparative Review of VAEs, GANs, Normalizing Flows, Energy-Based and Autoregressive Models. IEEE Trans. Pattern Anal. Mach. Intel., 1–1, doi:10.1109/TPAMI.2021.3116668 (2021).

20 Ramesh, A. et al. Zero-Shot Text-to-Image Generation in: International Conference on Machine Learning. 8821–8831 (PMLR, 2021). <https://proceedings.mlr.press/v139/ramesh21a.html>.

21 Oord, A. et al. Parallel WaveNet: Fast High-Fidelity Speech Synthesis in: International Conference on Machine Learning. 3918–3926 (PMLR, 2018). <https://proceedings.mlr.press/v80/oord18a.html>.

22 Noé, F., Tkatchenko, A., Müller, K.-R. & Clementi, C. Machine Learning for Molecular Simulation. Annu. Rev. Phys. Chem. 71, 361–390, doi:10.1146/annurev-physchem-042018-052331 (2020).

23 Noé, F. Machine Learning for Molecular Dynamics on Long Timescales in: Machine Learning Meets Quantum Physics (eds Kristof T. Schütt et al.) 331–372 (Springer International Publishing, 2020).

24 Xu, M., Luo, S., Bengio, Y., Peng, J. & Tang, J. Learning Neural Generative Dynamics for Molecular Conformation Generation in: International Conference on Learning Representations. (2021). <https://openreview.net/forum?id=pAbm1qfheGk>.

25 Satorras, V. G., Hoogeboom, E., Fuchs, F. B., Posner, I. & Welling, M. E(n) Equivariant Normalizing Flows. Adv. Neural Inf. Process. Syst. 34, 4181–4192 (2021).

26 Simm, G. & Hernandez-Lobato, J. M. A Generative Model for Molecular Distance Geometry in: International Conference on Machine Learning. 8949–8958 (PMLR, 2020). <https://proceedings.mlr.press/v119/simm20a.html>.

27 Noé, F., Olsson, S., Köhler, J. & Wu, H. Boltzmann generators: Sampling equilibrium states of many-body systems with deep learning. Science 365, doi:10.1126/science.aaw1147 (2019).

28 Fawzi, N. L., Parekh, S. H. & Mittal, J. Biophysical studies of phase separation integrating experimental and computational methods. Curr. Opin. Struct. Biol. 70, 78–86 (2021).

29 Goodfellow, I. et al. Generative Adversarial Nets. Adv. Neural Inf. Process. Syst. 27 (2014).

30 Woodard, J. et al. Intramolecular Diffusion in α-Synuclein: It Depends on How You Measure It. Biophys. J. 115, 1190–1199, doi:10.1016/j.bpj.2018.08.023 (2018).

31 Vaswani, A. et al. Attention is all you need. Adv. Neural Inf. Process. Syst. 30 (2017).

32 Battaglia, P. W. et al. Relational inductive biases, deep learning, and graph networks. arXiv preprint 1806.01261 (2018).

33 Wang, T.-C. et al. High-resolution image synthesis and semantic manipulation with conditional gans in: IEEE Conference on Computer Vision and Pattern Recognition. 8798–8807 (2018).

34 Durugkar, I., Gemp, I. & Mahadevan, S. Generative multi-adversarial networks. arXiv preprint 1611.01673 (2016).

35 Quaglia, F. et al. DisProt in 2022: improved quality and accessibility of protein intrinsic disorder annotation. Nucleic Acids Res. 50, D480–D487, doi:10.1093/nar/gkab1082 (2022).

36 Nguyen, H., Case, D. A. & Rose, A. S. NGLview-interactive molecular graphics for Jupyter notebooks. Bioinformatics (Oxford, England) 34, 1241–1242, doi:10.1093/bioinformatics/btx789 (2018).

37 Rosenbaum, D. et al. Inferring a continuous distribution of atom coordinates from cryo-EM images using VAEs. arXiv preprint 2106.14108 (2021).

38 Noé, F. & Rosta, E. Markov models of molecular kinetics. J. Chem. Phys. 151, 190401 (2019).

39 Miyato, T., Kataoka, T., Koyama, M. & Yoshida, Y. Spectral Normalization for Generative Adversarial Networks in: International Conference on Learning Representations. (2018). <https://openreview.net/forum?id=B1QRgziT->.

40 Esser, P., Rombach, R. & Ommer, B. Taming transformers for high-resolution image synthesis in: IEEE/CVF Conference on Computer Vision and Pattern Recognition. 12873–12883.

41 Dhariwal, P. & Nichol, A. Diffusion models beat gans on image synthesis. Adv. Neural Inf. Process. Syst. 34 (2021).

42 Eddy, S. R. Accelerated Profile HMM Searches. Plos Comp. Biol. 7, e1002195, doi:10.1371/journal.pcbi.1002195 (2011).

43 Dutagaci, B. et al. Charge-driven condensation of RNA and proteins suggests broad role of phase separation in cytoplasmic environments. eLife 10, e64004, doi:10.7554/eLife.64004 (2021).

44 Eastman, P. et al. OpenMM 7: Rapid development of high performance algorithms for molecular dynamics. Plos Comp. Biol. 13, e1005659, doi:10.1371/journal.pcbi.1005659 (2017).

45 Feig, M., Karanicolas, J. & Brooks, C. L. MMTSB Tool Set: enhanced sampling and multiscale modeling methods for applications in structural biology. J Mol Graph Model 22, 377–395, doi:10.1016/j.jmgm.2003.12.005 (2004).

46 Brooks, B. R. et al. CHARMM: The Biomolecular Simulation Program. J. Comput. Chem. 30, 1545–1614 (2009).

47 Goodfellow, I. Nips 2016 tutorial: Generative adversarial networks. arXiv preprint 1701.00160 (2016).

48 Isola, P., Zhu, J.-Y., Zhou, T. & Efros, A. A. Image-to-Image Translation with Conditional Adversarial Networks in: IEEE Conference on Computer Vision and Pattern Recognition (CVPR). 5967–5976 (2017).

49 Che, T. et al. Your GAN is secretly an energy-based model and you should use discriminator driven latent sampling. Adv. Neural Inf. Process. Syst. 33, 12275–12287 (2020).

50 Paszke, A. et al. PyTorch: An Imperative Style, High-Performance Deep Learning Library. Adv. Neural Inf. Process. Syst. 32, 8024–8035 (2019).

51 Kingma, D. P. & Ba, J. Adam: A method for stochastic optimization in: International Conference for Learning Representations. (2015). <https://openreview.net/forum?id=8gmWwjFyLj>.

52 Greener, J. G., Kandathil, S. M., Moffat, L. & Jones, D. T. A guide to machine learning for biologists. Nat. Rev. Mol. Cell Biol. 23, 40–55, doi:10.1038/s41580-021-00407-0 (2022).

53 Gulrajani, I., Ahmed, F., Arjovsky, M., Dumoulin, V. & Courville, A. C. Improved training of wasserstein gans. Adv. Neural Inf. Process. Syst. 30 (2017).

