## supplementary figures and tables for "Direct Generation of Protein Conformational Ensembles via Machine Learning"

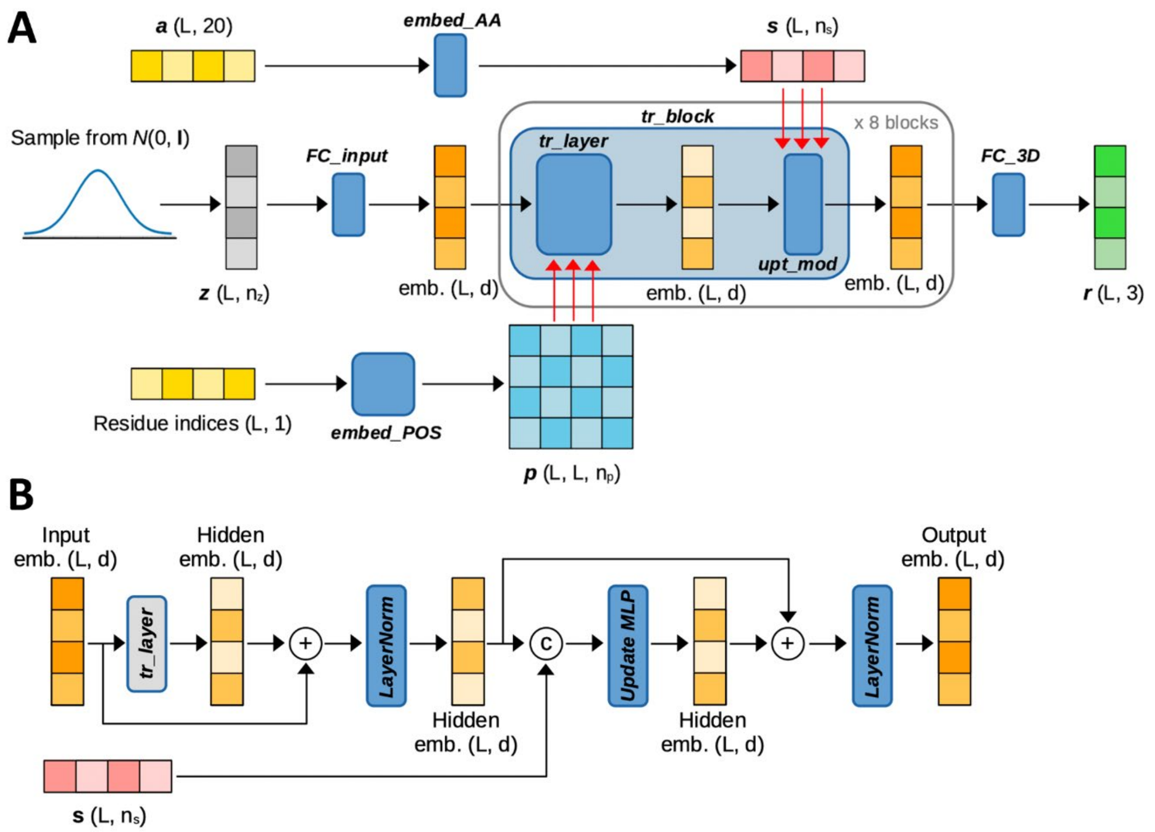


**Supplementary Fig. 1. Architecture of the generator network of idpGAN.** For a protein of length $L$, a latent tensor $\boldsymbol{z} \in \mathbb{R}^{L \times n_z}$, with $n_z$ = 16, is sampled from a Gaussian prior. This tensor is converted in an embedding sequence of $L$ tokens with dimension $d$ = 64 by a fully-connected module acting position-wise (*FC_input*, see **Supplementary Table 2**). The embedding is then processed by a series of $n_t$= 8 transformer blocks (*tr_block*). Each block is composed of two sub-modules, a "transformer layer" (*tr_layer*) and an "updater module" (*upt_mod*) based on a previous network (see ref. 31 in the main text). A "transformer layer" receives as input an embedding and updates it through a self-attention mechanism (ref. 31 in the main text). We use $n_h$ = 8 attention heads and $d_{model}$ = 16 as hyper-parameters. Different from the original transformer, each layer uses 2D relative position encoding $\boldsymbol{p} \in \mathbb{R}^{L \times L \times n_p}$ (with $n_p$ = 64) as in AF2 (ref. 10 in the main text). The encoding $\boldsymbol{p}$ is derived through an embedding module (*embed_POS*), where each pair of residues in a protein is labeled with a learnable $n_p$-dimensional vector associated with their sequence separation (i.e., the difference between residue indices). The separation values are clipped from -24 to 24. The same $\boldsymbol{p}$ is used in all blocks of the network (triple red arrows in the figure). For each block, we first linearly project $\boldsymbol{p}$ onto a bias term for each attention head and then add the biases to the logits values of the corresponding attention maps. The "updater modules", shown in panel **B**, are composed of two layer normalization operations and a fully-connected module (*update_MLP*, see **Supplementary Table 2**). In contrast to (29), we do not use dropout regularization, since we found it to negatively impact our GAN performance. An "updater module" also receives as input an encoding $\boldsymbol{s} \in \mathbb{R}^{L \times n_s}$ (with $n_s$ = 32). In $\boldsymbol{s}$, each of the 20 amino acid types is associated with a learnable $n_s$-dimensional vector through an embedding layer (*embed_AA*) that takes as input a tensor $\boldsymbol{a} \in \mathbb{R}^{L \times 20}$ storing one hot encodings for amino acid types. To inject $\boldsymbol{s}$, we concatenate it along the feature axis to the tensor produced by the first layer normalization. The same $\boldsymbol{s}$ is injected into all blocks of the network (triple red arrows in the figure). The output of the final block is converted in a molecular conformation $\boldsymbol{r}$ through a fully-connected module acting position-wise (*FC_3D*, see **Supplementary Table 2**).

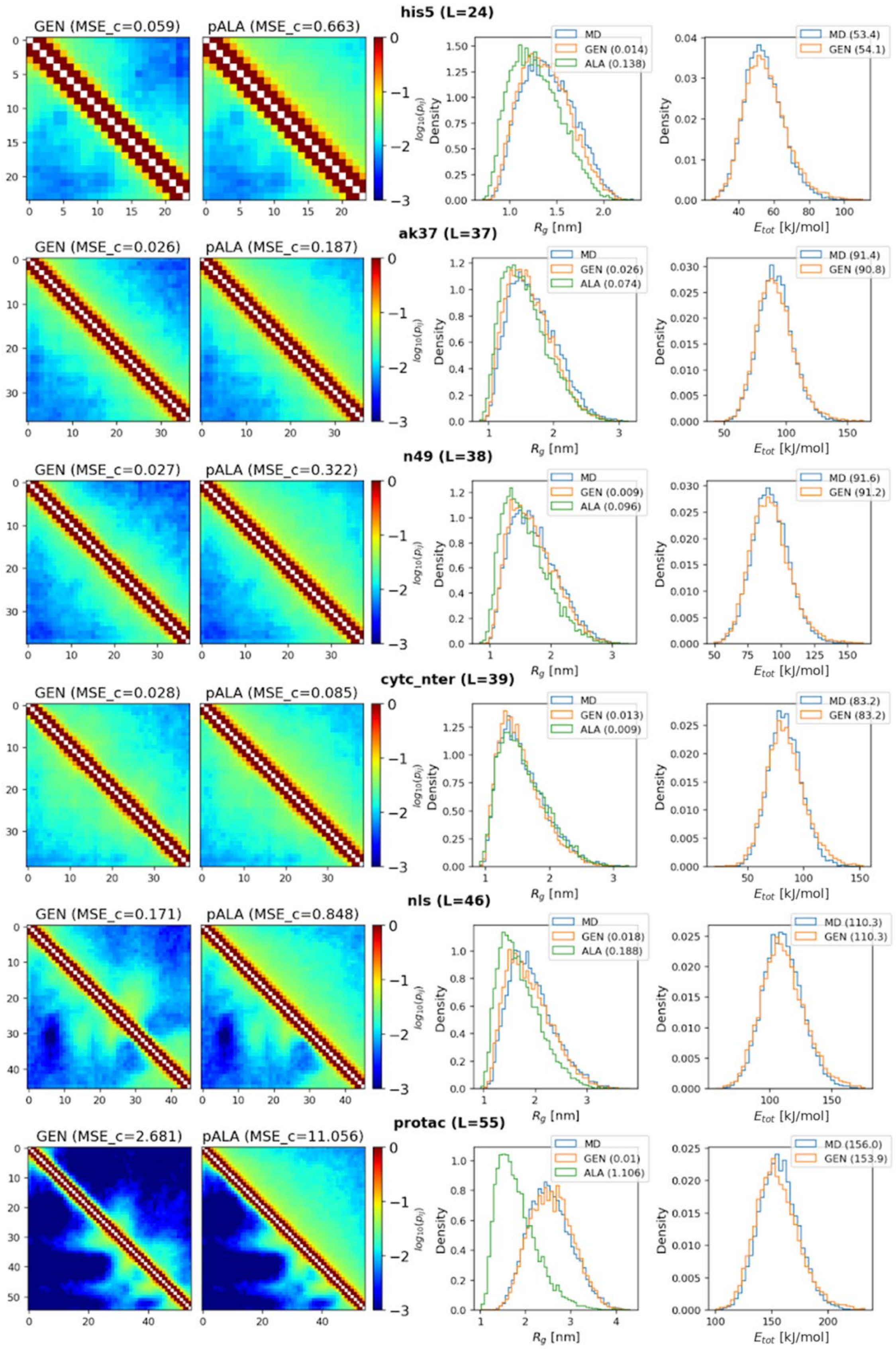


**Supplementary Fig. 2. Evaluation of idpGAN on *IDP_test* proteins.** Contact maps, radius-of-gyration distributions, and energy distributions from generated (GEN) and polyAla (pALA) ensembles vs. ensembles from MD snapshots for *his5*, *ak37*, *n49*, *cytc_nter*, *nls*, and *protac*. See **Fig. 2** for more details.

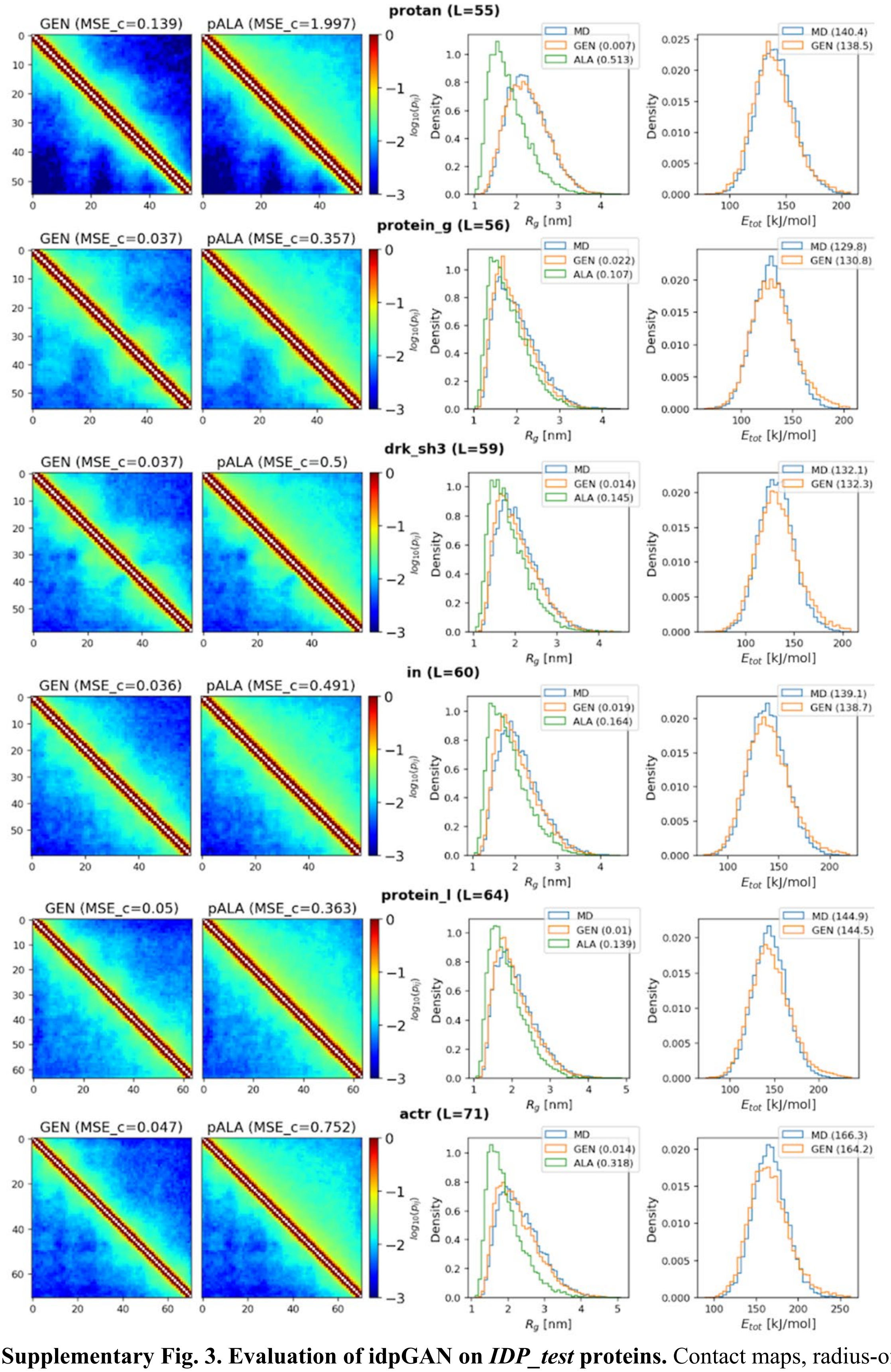


**Supplementary Fig. 3. Evaluation of idpGAN on *IDP_test* proteins.** Contact maps, radius-of-gyration distributions, and energy distributions from generated (GEN) and polyAla (pALA) ensembles vs. ensembles from MD snapshots for *protan*, *protein_g*, *drk_sh3*, *in*, *protein_l*, and *actr*. See **Fig. 2** for more details.

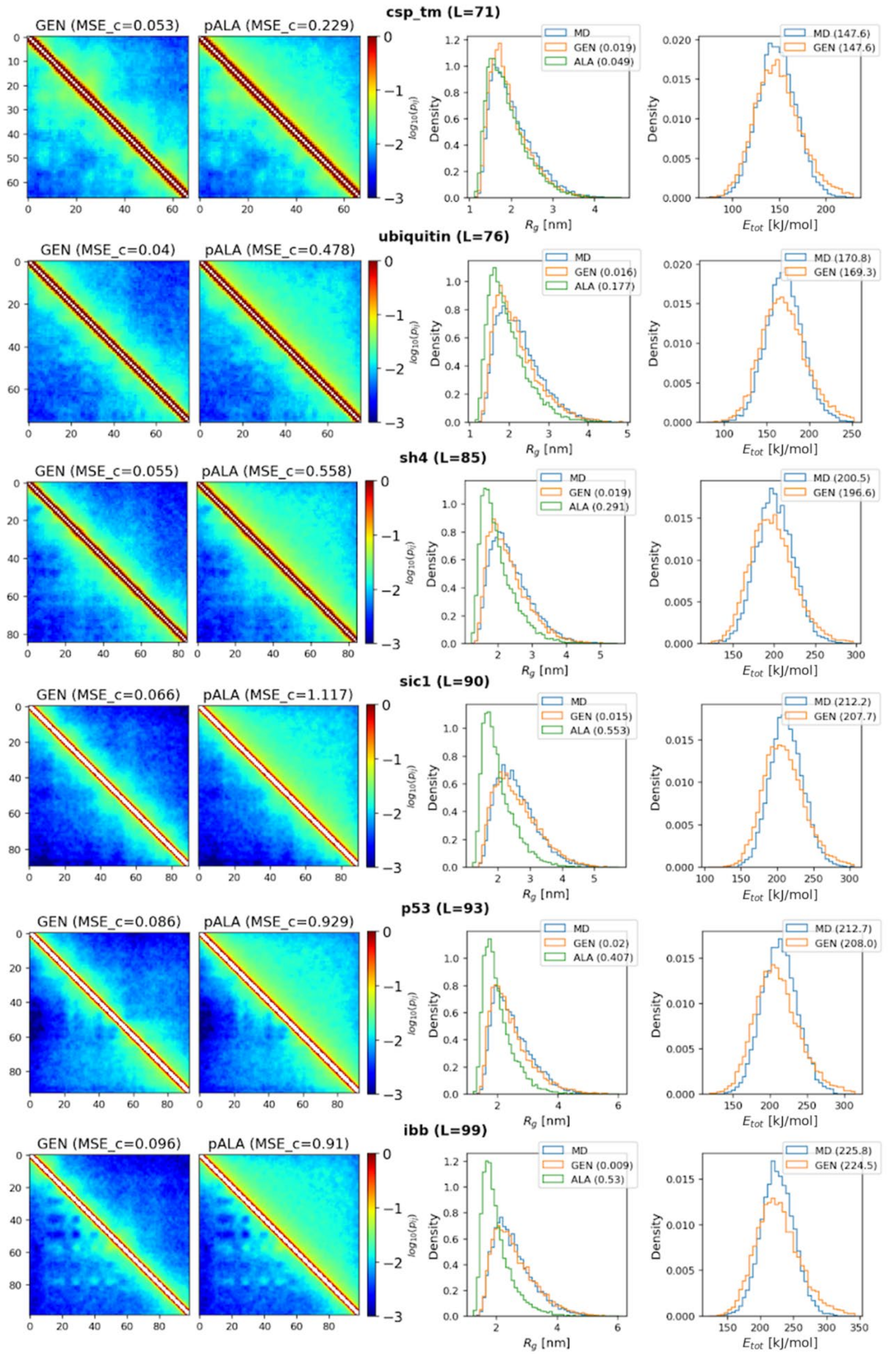


**Supplementary Fig. 4. Evaluation of idpGAN on *IDP_test* proteins.** Contact maps, radius-of-gyration distributions, and energy distributions from generated (GEN) and polyAla (pALA) ensembles vs. ensembles from MD snapshots for *csp_tm*, *ubiquitin*, *sh4*, *sic1*, *p53*, and *ibb*. See **Fig. 2** for more details.

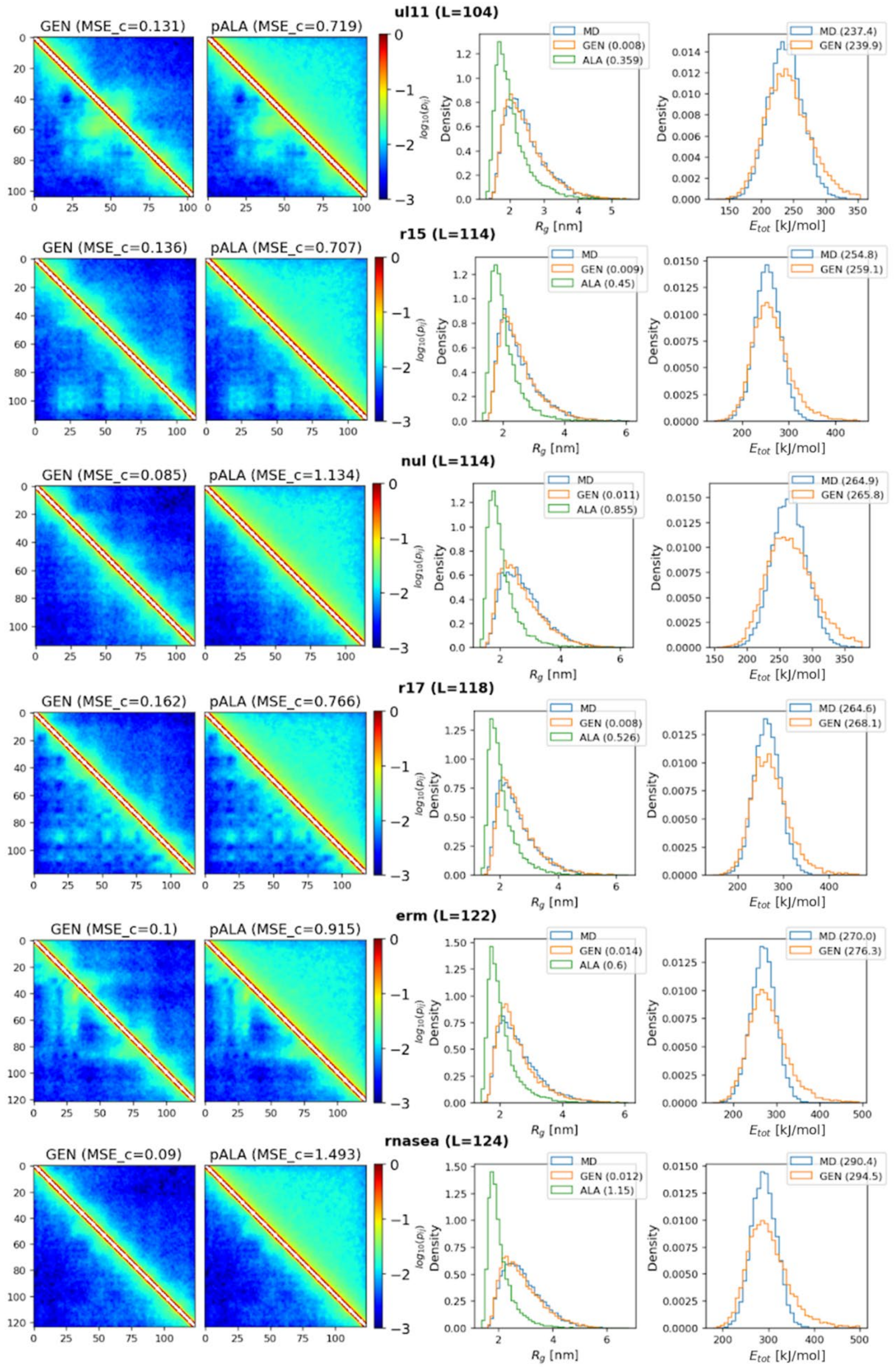


**Supplementary Fig. 5. Evaluation of idpGAN on *IDP_test* proteins.** Contact maps, radius-of-gyration distributions, and energy distributions from generated (GEN) and polyAla (pALA) ensembles vs. ensembles from MD snapshots for *ul11*, *r15*, *nul*, *r17*, *erm*, and *rnasea*. See **Fig. 2** for more details.

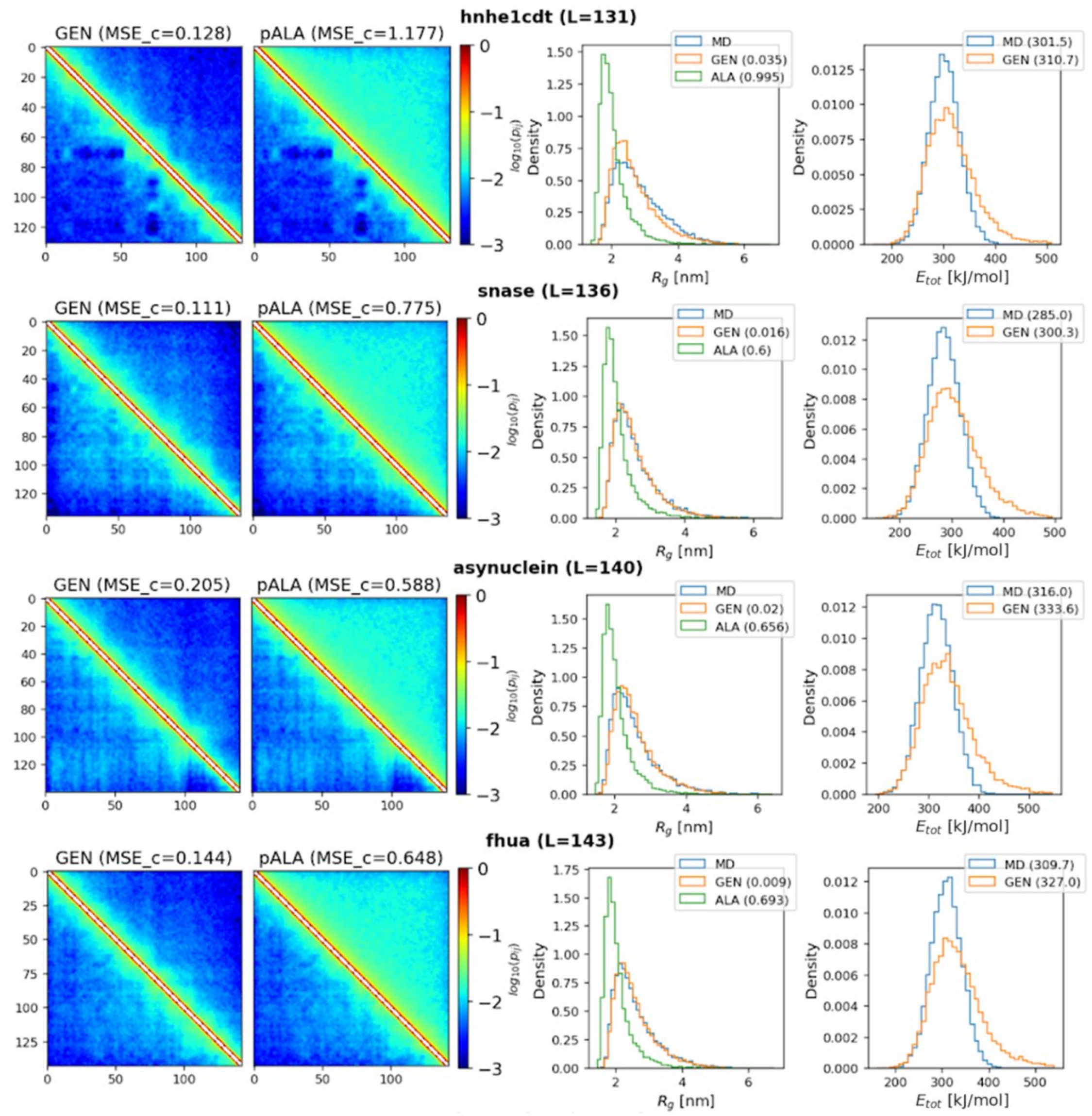


**Supplementary Fig. 6. Evaluation of idpGAN on *IDP_test* proteins.** Contact maps, radius-of-gyration distributions, and energy distributions from generated (GEN) and polyAla (pALA) ensembles vs. ensembles from MD snapshots for *hnhe1cdt*, *snase*, *asynuclein*, and *fhua*. See **Fig. 2** for more details.

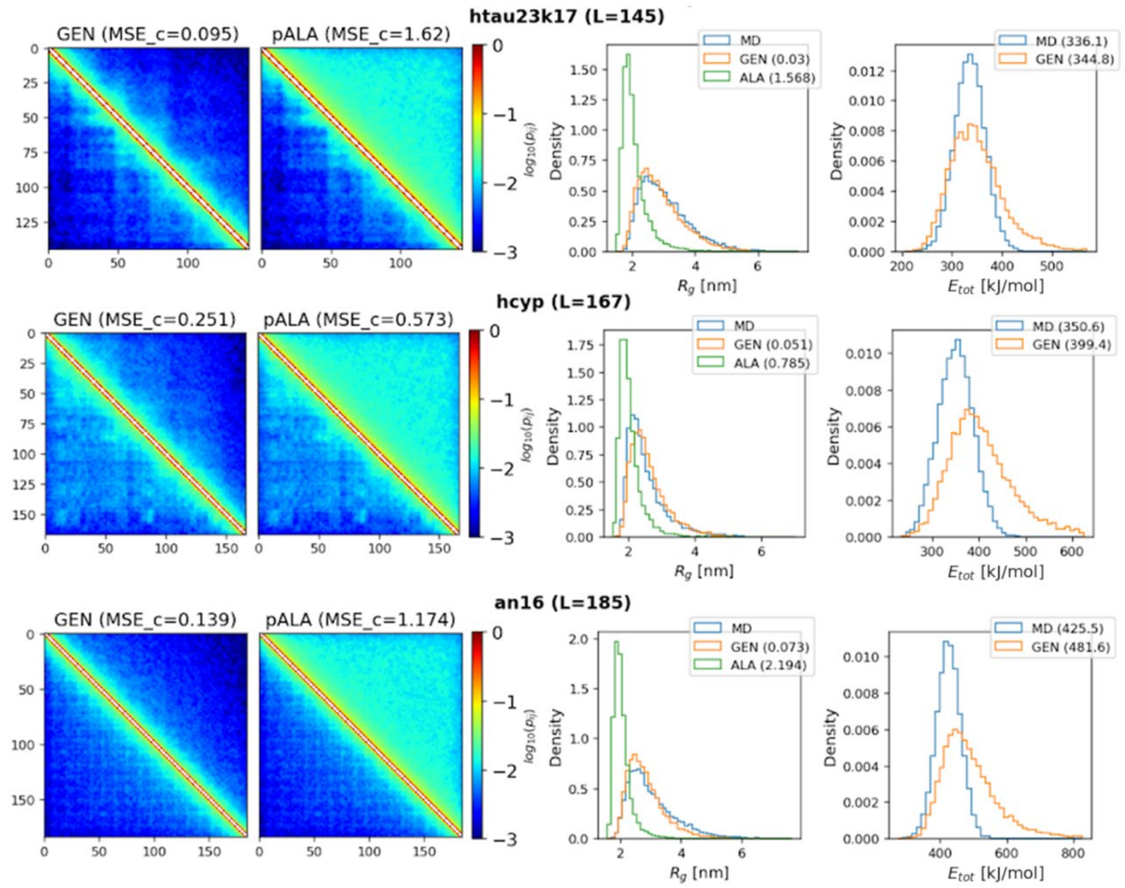


**Supplementary Fig. 7. Evaluation of idpGAN on *IDP_test* proteins.** Contact maps, radius-of-gyration distributions, and energy distributions from generated (GEN) and polyAla (pALA) ensembles vs. ensembles from MD snapshots for *htau23k17*, *hcyp*, and *an16*. See **Fig. 2** for more details

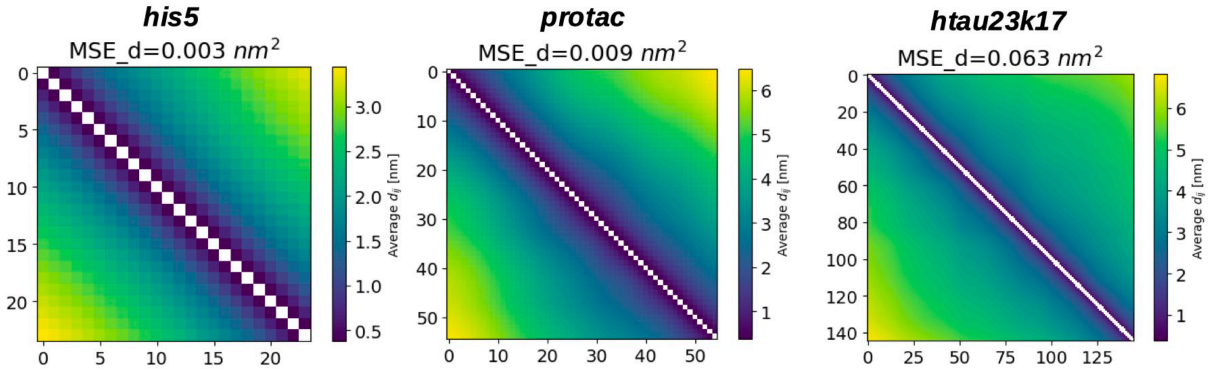


**Supplementary Fig. 8. Average distance maps.** The figures show the average distance maps for ensembles from idpgGAN (upper triangle of the images) and MD simulations (lower triangle) for the *his5*, *protac* and *htau23k17* proteins of the *IDP_test* set. The *MSE_d* scores of the idpGAN maps with the MD ones are shown above each figure.

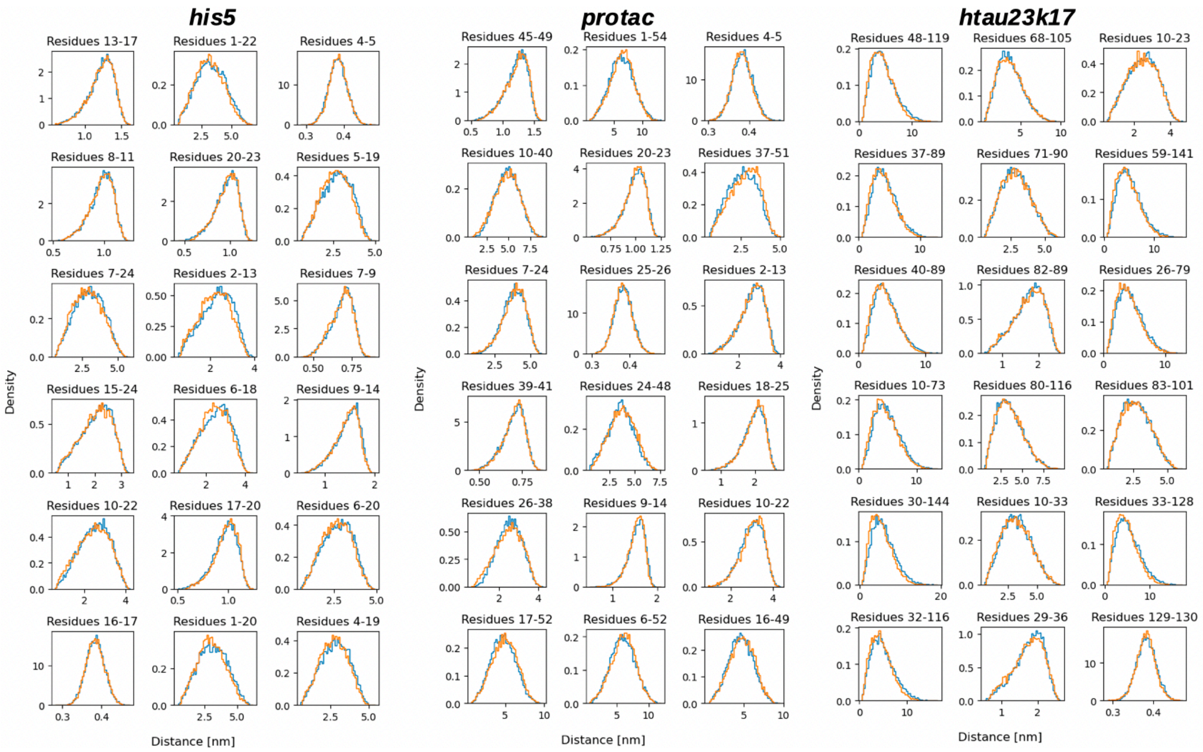


**Supplementary Fig. 9. Histograms of Cα-Cα distances.** The figures show distance distributions for the MD (blue color) and idpGAN (orange) ensembles for the *his5*, *protac* and *htau23k17* proteins of the *IDP_test* set. For each protein, we show randomly selected distance distributions with the indices of the residues shown above each histogram.

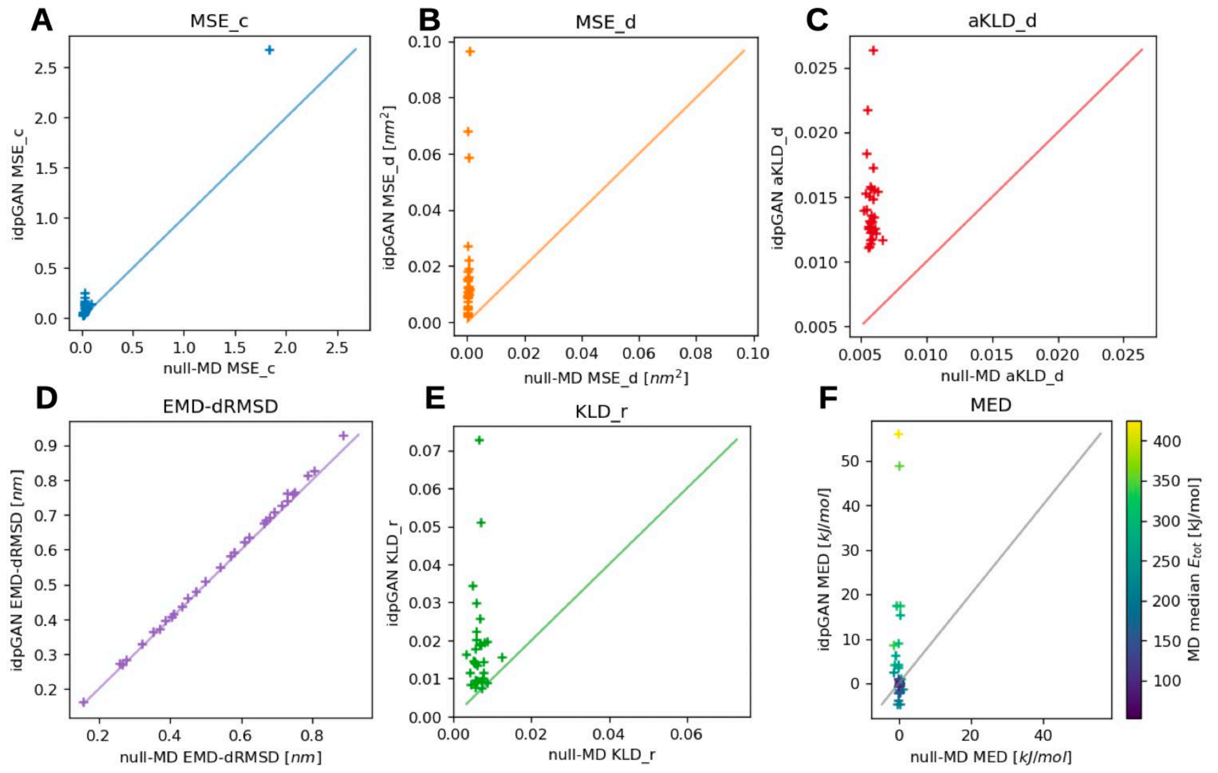


**Supplementary Fig. 10. Evaluation of idpGAN and null-MD ensembles for approximating reference MD data.** Results are reported for *IDP_test* set protein ($N = 31$). **A**, **B**, **C**, **D**, and **E** show the values of *MSE_c*, *MSE_d*, *aKLD_d*, *EMD-dRMSD* and *KLD_r*, respectively, obtained by extracting snapshots from a long independent MD simulation (null-MD) (x-axis) and idpGAN (y-axis) for all the proteins in the set. Lower values indicate a better performance in approximating reference MD ensembles (obtained from 5 shorter MD simulations). *MED* values of idpGAN and null-MD ensembles are confronted in **F** with markers colored according to the median potential energy of proteins in the reference MD ensembles.

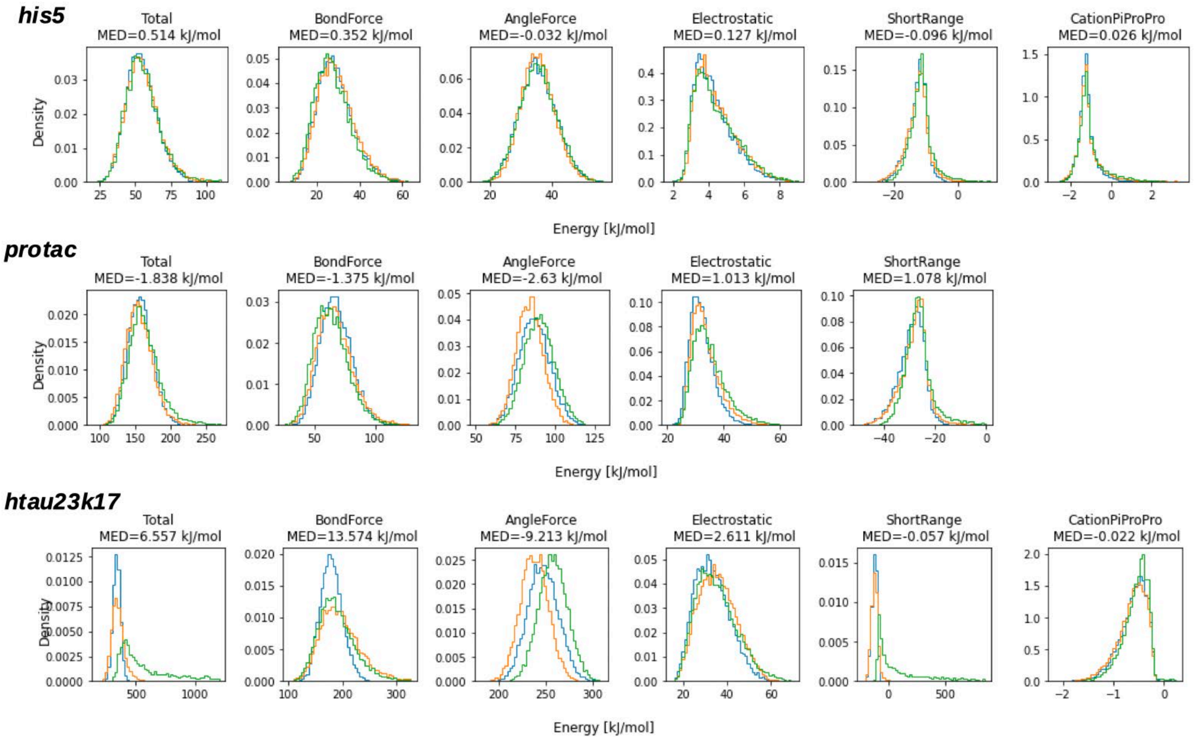


**Supplementary Fig. 11. Histograms of potential energy terms for the *his5*, *protac* and *htau23k17* proteins of the *IDP_test*.** The histograms show data for the ensembles from MD (blue color), idpGAN (orange) and idpGAN trained without the additional loss term for removing steric clashes (green). The name of each energy term is shown on the top of the histograms, along with the difference in median potential energy values between the MD and idpGAN ensembles for each term. Note that for *protac* the cation-π term is missing, since it does not have any aromatic residues.

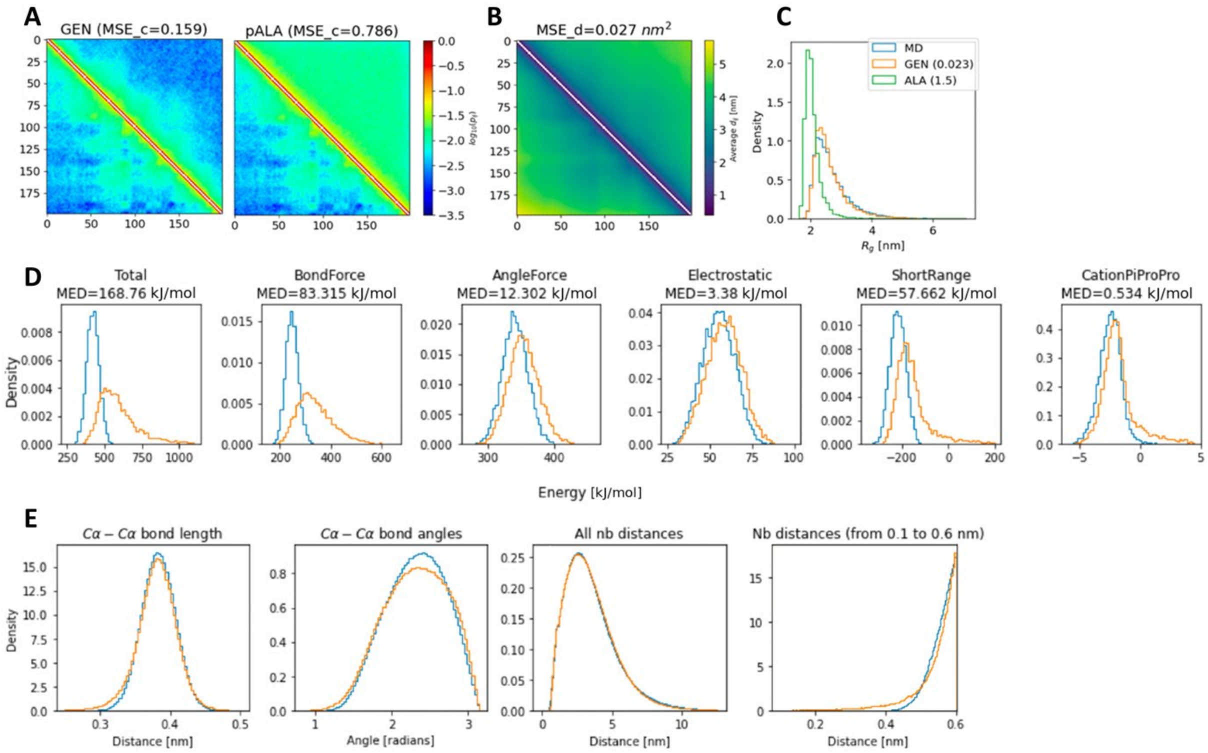


**Supplementary Fig. 12. Generated ensemble of *DP02478r001* (*L* = 199), an IDP of the *HB_val* set. a** Contact maps of the idpGAN and polyAla ensembles (upper triangles) compared to the MD one (lower triangles). **b** Average distance map of the MD (lower triangle) and idpGAN (upper triangle) ensembles. **C** Distributions of radius of gyration for the MD, idpGAN and polyAla ensembles. Please refer to **Fig. 2** in the main text for more information on panel **A** and **C**, and to **Supplementary Fig. 8** for panel **B**. **D** Histograms showing values of potential energy terms for the MD (blue color) and idpGAN (orange) ensembles. The difference in median potential energy between the idpGAN and MD ensembles is shown for each term. **E** Histograms showing the distribution of Cα-Cα bond lengths, bond angles and non-bonded (“nb”) distances. These are the geometrical features that determine the values of the “BondForce”, “AngleForce” and “ShortRange” terms respectively. The non-bonded distances histogram on the left shows all distances from 0 to 12 nm. The one on the right zooms in the 0.1 to 0.6 nm range (shaded yellow area in the left histogram), which contains short distances that cause high energy values for the “ShortRange” term. The overall distribution of distances is captured by idpGAN, while there are small divergences in the 0.1 to 0.6 nm range.

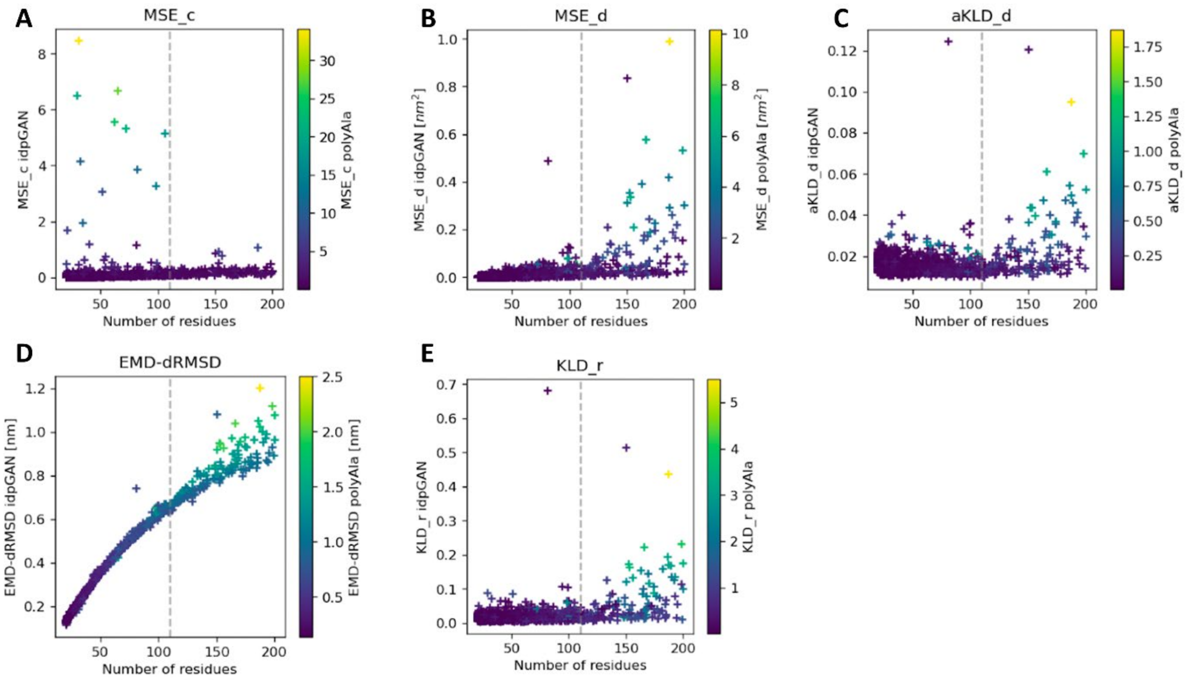


**Supplementary Fig. 13. Evaluation of idpGAN ensembles for approximating MD ensembles.** The panels from **A** to **E** show the values of *MSE_c*, *MSE_d*, *aKLD_d*, *EMD-dRMSD*, and *KLD_r* metrics respectively obtained by idpGAN for IDPs of the *HB_val* set as a function of protein length. The markers are colored according to the scores obtained with each metric by the polyAla approximation strategy. The dashed vertical lines represent the maximum crop length used in idpGAN training ($L$ = 110). The scores of most evaluation metrics do not show a strong dependence on IDP length, with the exception of *EMD-dRMSD*. Note that this dependence arises naturally for *EMD-dRMSD*, since we use a fixed number of conformations $n_{eval}$ = 10,000 to evaluate each IDP. When calculating *EMD-dRMSD*, each reference conformation is paired to a generated one to minimize a global dRMSD score. Since the size of the conformational space of an IDP increases with length, the expected dRMSD value between a reference conformation and the most similar one in a generated ensemble also tends to increase with IDP length if $n_{eval}$ (the number of conformations in the generated ensemble) is kept constant.

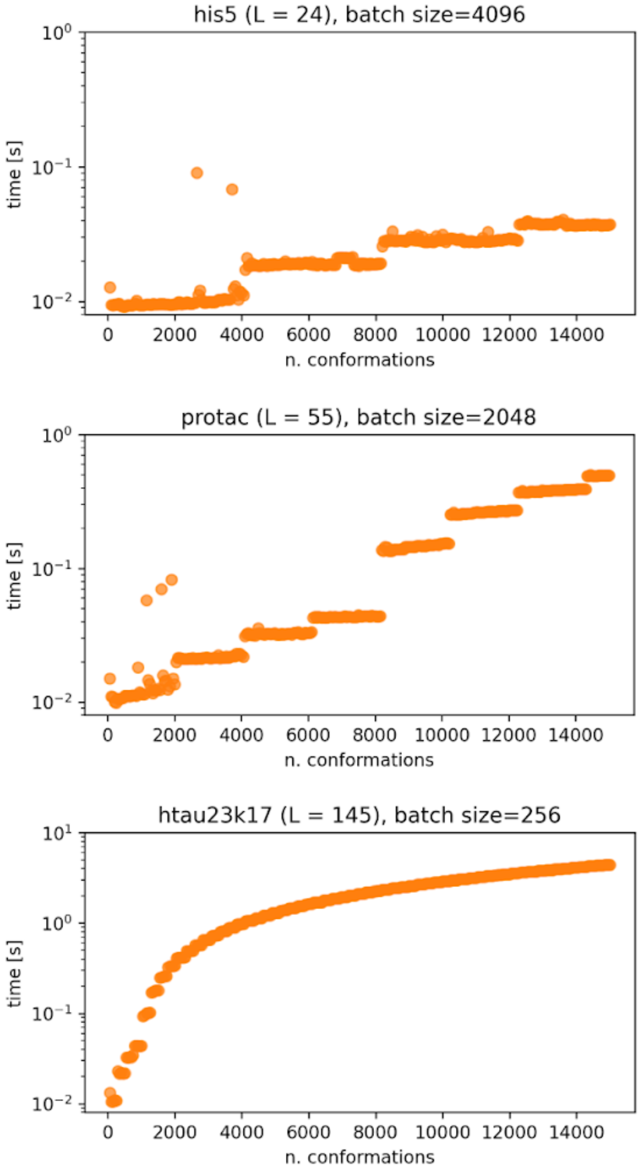


**Supplementary Fig. 14. GPU time used by the G network to generate a conformational ensemble as a function of the number of conformations in the ensemble.** We show data for the *his5*, *protac* and *htau23k17* proteins of the *IDP_test* set. Beside the name of the protein, its number of amino acids *L* and the batch size used to generate all the samples in the ensembles are shown. Please refer to the “Methods” section in the main text and **Supplementary Table 3** for details.

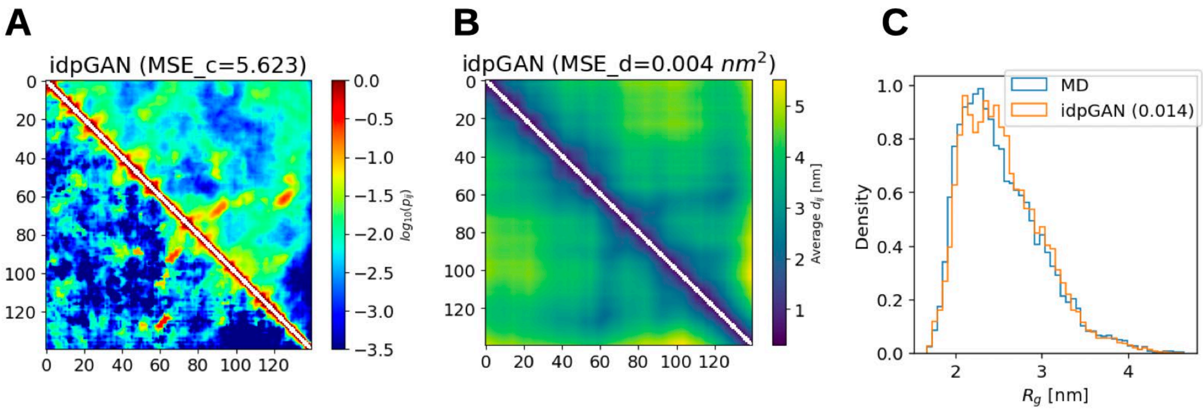


**Supplementary Fig. 15 Comparison of idpGAN ensembles for α-synuclein with the training data ensemble.** **A** idpGAN contact map (in the upper triangle of the image) confronted with the MD map from the training data of the model (lower triangle). Their *MSE_c* score is shown in brackets. **B** idpGAN average distance map (upper triangle) confronted with the corresponding MD map from the training data (lower triangle). Their *MSE_d* score is shown in brackets. **C** Radius-of-gyration distributions for the training MD data (blue) and idpGAN (orange) ensembles with the *KLD_r* value shown in brackets.

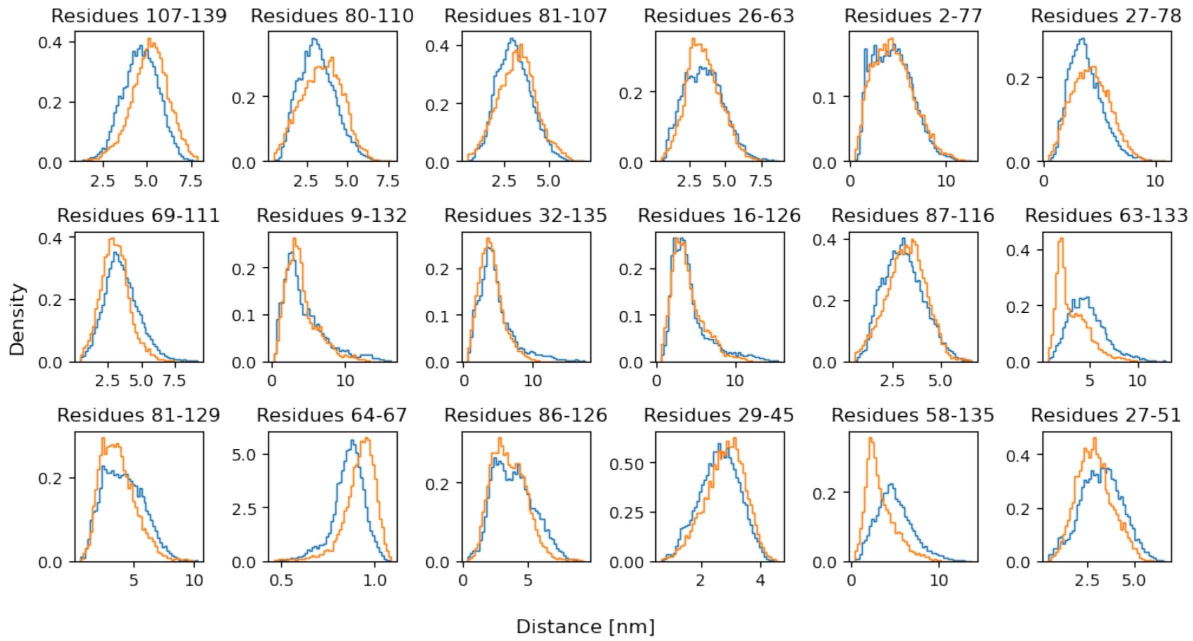


**Supplementary Fig. 16. Histograms for Cα-Cα distances for all-atom α-synuclein conformations.** Data for the validation MD (blue color) and idpGAN (orange) ensembles is reported. 18 distance distributions between randomly selected residues are shown (the indices of the residues are indicated above each histogram).

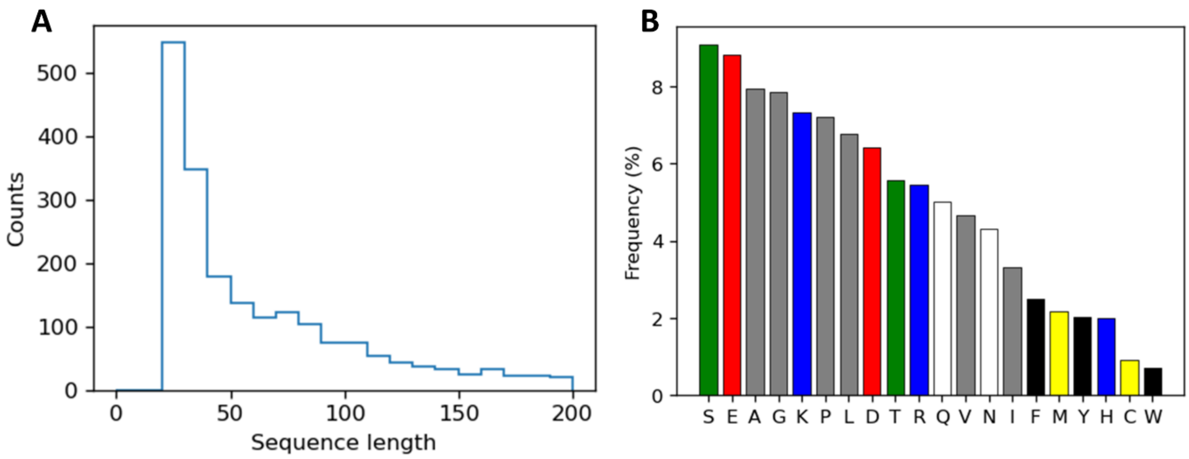


**Supplementary Fig. 17.** Properties of the 1966 IDP sequences from DisProt IDRs in the training set of idpGAN. **A:** Histogram of the sequence lengths of the IDPs. **B:** Amino acid frequencies in the IDRs. The amino acids are colored according to their physiochemical characteristics: green is for "small hydroxy", red is for "acidic", gray is for "aliphatic", blue is for "basic", white is for "amide", black is for "aromatic", yellow is for "sulfur". Colors were adopted from a similar plot in https://www.ebi.ac.uk/uniprot/TrEMBLstats showing amino acid frequencies in the whole TrEMBL database.

**Supplementary Table 1.** Sequences of the proteins in the *IDP_test* set.

| Name | Len. | $R_g$ (nm) | Sequence | Ref. |
|---|---|---|---|---|
| his5 | 24 | 1.38 | DSHAKRHHGYKRKFHEKHHSHRGY | 1 |
| ak37 | 37 | 1.69 | AAKAAAAKAAAAKAAAAKAAAAKAAAAKAAAAKAAGY | 2 |
| n49 | 38 | 1.37 | GCQTSRGLFGNNNTNNINNSSSGMNNASAGLFGSKPFA | 3 |
| cytc_nter | 39 | 1.84 | MIFFMVMPIMIGGFGNWLVPLMIGAPDMAFPRMNNSFWL | 2 |
| nls | 46 | 1.63 | ACETNKRKREQISTDNEAKMQIQEEKSPKKKRKKRSSKANKPPEFA | 3 |
| protac | 55 | 3.00 | CEEGGEEEEEEEEGDGEEEDGDEDEEAESATGKRAAEDDEDDDVDTKKQKTDEDC | 4 |
| protan | 55 | 2.55 | CDAAVDTSSEITTKDLKEKKEVVEEAENGRDAPANGNANEENGEQEADNEVDEEC | 4 |
| protein_g | 56 | 2.30 | MQYKLALNGKTLKGETTTEAVDAATAEKVFKQYANDNGVDGEWAYDDATKTFAVTE | 5 |
| drk_sh3 | 59 | 2.19 | MEAIAKHDFSATADDELSFRKTQILKILNMEDDSNWYRAELDGKEGLIPSNYIEMKNHD | 6 |
| in | 60 | 2.16 | GSHCFLDGIDKAQEEHEKYHSNWRAMASDFNLPPVVAKEIVASCDKCQLKGEAMHGQVDC | 7 |
| protein_l | 64 | 1.65 | MEEVTIKANLIFANGSTQTAEFKGTFEKATSEAYAYADTLKKDNGEWTVDVADKGYTLNIKFAG | 8 |
| actr | 71 | 2.51 | GTQNRPLLRNSLDDLVGPPSNLEGQSDERALLDQLHTLLSNTDATGLEEIDRALGIPELVNQGQALEPKQD | 9 |
| csp_tm | 67 | 1.47 | GPGMCRGKVKFFDSKKGYGFITKDEGGDVFVHFSAIEMEGFKTLKEGQVVEFEIQEGKKGGQAAHVKVVEC | 4 |
| ubiquitin | 76 | 2.52 | MQIFVKTLTGKTITLEVEPSDTIENVKAKIQDKEGIPPDQQRLIWAGKQLEDGRTLSDYNIQKESTLHLVLRLRGG | 2 |
| sh4 | 85 | 2.82 | MGSNKSKPKDASQRRRSLEPAENVHGAGGGAFPASQTPSKPASADGHRGPSAAFAPAAAEPKLFGGFNSSDTVTSPQRAGPLAGG | 10 |
| sic1 | 90 | 3.00 | MTPSTPPRSRGTRYLAQPSGNTSSSALMQGQKTPQKPSQNLVPVTPSTTKSFKNAPLLAPPNSNMGMTSPFNGLTSPQRSPFPKSSVKRT | 11 |
| p53 | 93 | 2.87 | MEEPQSDPSVEPPLSQETFSDLWKLLPENNVLSPLPSQAMDDLMLSPDDIEQWFTEDPGPDEAPRMPEAAPPVAPAPAAPTPAAPAPAPSWPL | 12 |
| ibb | 99 | 2.52 | GCTNENANTPAARLHRFKNKGKDSTEMRRRRIEVNVELRKAKKDDQMLKRRNVSSFPDDATSPLQENRNNQGTVNWSVDDIVKGINSSNVENQLQATFA | 3 |
| ul11 | 104 | 2.43 | MGLSFSGTRPCCCRNNVLITDDGEVVSLTAHDFDVVDIESEEEGNFYVPPDMRGVTRAPGRQRLRSSDPPSRHTHRRTPGGACPATQFPPPMSDSEWSHPQFEK | 13 |
| r15 | 114 | 2.33 | KLKEACKQQNFNTGIKDFDFWLSEVEALLASEDYGKDLASVNNLLKKHQLLEADISAHEDRLKDLNSQADSLMTSSAFDTSQVKDKRETINGRFQRIKCMAAARRAKLNESHRL | 7 |
| nul | 114 | 2.66 | GCGFKGFDTSSSSSNSAASSSFKFGVSSSSSGPSQTLTSTGNFKFGDQGGFKIGVSSDSGSINPMSEGFKFSKPIGDFKFGVSSESKPEEVKKDSKNDNFKFGLSSGLSNPVFA | 3 |
| r17 | 118 | 2.37 | GSRLEESCEYQQFVANVEEEEAWINEKMTLVASEDYGDTLAAIQGLLKKHEAFETDFTVHKDRVNDVAANGEDLIKKNNHHVENITAKMKGLKGKVSDLECAAAQRKAKLDENSAFLQ | 7 |
| erm | 122 | 3.96 | MDGFYDQQVPFMVPGKSRSEECRGRPVIDRKRKFLDTDLAHDSEELFQDLSQLQEAWLAEAQVPDDEQFVPDFQSDNLVLHAPPPTKIKRELHSPSSELSSCSHEQALGANYGEKCLYNYCA | 14 |
| rnasea | 124 | 3.36 | KETAAAKFERQHMDSSTSAASSSNYCNQMMKSRNLTKDRCKPVNTFVHESLADVQAVCSQKNVACKNGQTNCYQSYSTMSITDCRETGSSKYPNCAYKTTQANKHIIVACEGNPYVPVHFDASV | 15 |
| hnhe1cdt | 131 | 3.63 | MVPAHKLDSPTMSRARIGSDPLAYEPKEDLPVITIDPASPQSPESVDLVNEELKGKVLGLSRDPAKVAEEDEDDDGGIMMRSKETSSPGTDDVFTPAPSDSPSSQRIQRCLSDPGPHPEPGEGEPFFPKGQ | 9 |
| snase | 136 | 2.12 | ATSTKKLHKEPATLIKAIDGDTVKLMYKGQPMTFRLLLVDTPETKHPKKGVEKYGPEASAFTKKMVENAKKIEVEFDKGQRTDKYGRGLAYIYADGKMVNEALVRQGLAKVAYVYKPNNTHEQHLRKSEAQAKKEK | 16 |
| asynuclein | 140 | 3.30 | MDVFMKGLSKAKEGVVAAAEKTKQGVAEAAGKTKEGVLYVGSKTKEGVVHGVATVAEKTKEQVTNVGGAVVTGVTAVAQKTVEGAGSIAAATGFVKKDQLGKNEEGAPQEGILEDMPVDPDNEAYEMPSEEGYQDYEPEA | 17 |
| fhua | 143 | 3.34 | ESAWGPAATIAARQSATGTKTDTPIQKVPQSISVVTAEEMALHQPKSVKEALSYTPGVSVGTRGASNTYDHLIIRGFAAEGQSQNNYLNGLKLQGNFYNDAVIDPYMLERAEIMRGPVSVLYGKSSPGGLLNMVSKRPTTEPL | 15 |
| htau23k17 | 145 | 3.60 | MSSPGSPGTPGSRSRTPSLPTPPTREPKKVAVVRTPPKSPSSAKSRLQTAPVPMPDLKNVKSKIGSTENLKHQPGGGKVQIVYKPVDLSKVTSKCGSLGNIHHKPGGGQVEVKSEKLDFKDRVQSKIGSLDNITHVPGGGNKKIE | 18 |
| hCyp | 167 | 2.51 | GPMCNPTVFFDIAVDGEPLGRVSFELFADKVPKTAENFRALSTGEKGFGYKGSSFHRIIPGFMSQGGDFTRHNGTGGKSIYGEKFEDENFILKHTGPGILSMANAGPNTNGSQFFISTAKTEFLDGKHVVFGKVKEGMNIVEAMERFGSRNGKTSKKITIADSGQLC | 7 |
| an16 | 185 | 4.44 | MHHHHHHPGAPAQTPSSQYGAPAQTPSSQYGAPAQTPSSQYGAPAQTPSSQYGAPAQTPSSQYGAPAQTPSSQYGAPAQTPSSQYGAPAQTPSSQYGAPAQTPSSQYGAPAQTPSSQYGAPAQTPSSQYGAPAQTPSSQYGAPAQTPSSQYGAPAQTPSSQYGAPAQTPSSQYGAPAQTPSSQYGAPAQTPSSQYV | 2 |

**Supplementary Table 2.** Architecture and hyper-parameters of the neural network modules.

| **Module** | **Neural network** | **Structure** |
|---|---|---|
| *FC_input* | Generator | *Input → {LinearLayer($n_{in}$=16, $n_{out}$=64) → LeakyReLU(0.2) → LinearLayer($n_{in}$=64, $n_{out}$=64) → LeakyReLU(0.2) → LinearLayer($n_{in}$=64, $n_{out}$=64)} → Output*. |
| *update_MLP* | Generator | *Input → {LinearLayer($n_{in}$=64+32, $n_{out}$=128) → LeakyReLU(0.2) → LinearLayer($n_{in}$=128, $n_{out}$=64)} → Output.* |
| *FC_3D* | Generator | *Input → {LinearLayer($n_{in}$=64, $n_{out}$=64) → LeakyReLU(0.2) → LinearLayer($n_{in}$=64, $n_{out}$=3)} → Output.* |
| MLP_discriminator | Discriminator for CG data | *Input → {LinearLayer($n_{in}$=L*(L-1)/2+20*L, $n_{out}$=1024, sn=True) → LeakyReLU(0.2) → LinearLayer($n_{in}$=1024, $n_{out}$=1024, sn=True) → LeakyReLU(0.2) → LinearLayer($n_{in}$=1024, $n_{out}$=1, sn=True) → Sigmoid()} → Output.* |
| MLP_discriminator | Discriminator for full-atom data | *Input → {LinearLayer($n_{in}$=L*(L-1)/2+(L-3), $n_{out}$=1024) → LeakyReLU(0.2) → LinearLayer($n_{in}$=1024, $n_{out}$=1024) → LeakyReLU(0.2) → LinearLayer($n_{in}$=1024, $n_{out}$=1)} → Output.* |

The *sn* argument in the linear layers indicates the use of spectral normalization (see main text).

**Supplementary Table 3.** Parameters used for generating conformations with idpGAN during timing tests.

| Protein length interval | Batch size |
|---|---|
| (0, 50] | 4096 |
| (50, 80] | 2048 |
| (80, 110] | 1024 |
| (110, 140] | 512 |
| (140, 200] | 256 |

**Supplementary Table 4.** Hyper-parameters in generator network of idpGAN for CG and all-atom Cα trace modeling.

| **Hyper-parameter name** | **CG data** | **All-atom data** |
|---|---|---|
| Transformer block embedding dimension ($d$) | 64 | 128 |
| Feed forward dimension in the updater module of transformer blocks | 128 | 256 |
| Number of transformer blocks ($n_t$) | 8 | 16 |
| Number of attention heads ($n_h$) | 8 | 12 |
| Dimension of the query, key and values vectors ($d_{model}$) | 16 | 32 |
| Maximum sequence separation in relative 2d positional embeddings | 24 | 32 |

**Supplementary References**


1 Cragnell, C., Durand, D., Cabane, B. & Skepo, M. Coarse-grained modeling of the intrinsically disordered protein Histatin 5 in solution: Monte Carlo simulations in combination with SAXS. *Proteins* **84**, 777-791, doi:10.1002/prot.25025 (2016).

2 Kohn, J. E. *et al.* Random-coil behavior and the dimensions of chemically unfolded proteins. *Proc Natl Acad Sci U S A* **101**, 12491-12496, doi:10.1073/pnas.0403643101 (2004).

3 Fuertes, G. *et al.* Decoupling of size and shape fluctuations in heteropolymeric sequences reconciles discrepancies in SAXS vs. FRET measurements. *Proc Natl Acad Sci U S A* **114**, E6342-E6351, doi:10.1073/pnas.1704692114 (2017).

4 Muller-Spath, S. *et al.* From the Cover: Charge interactions can dominate the dimensions of intrinsically disordered proteins. *Proc Natl Acad Sci U S A* **107**, 14609-14614, doi:10.1073/pnas.1001743107 (2010).

5 Smith, C. K. *et al.* Surface point mutations that significantly alter the structure and stability of a protein's denatured state. *Protein Sci* **5**, 2009-2019, doi:10.1002/pro.5560051007 (1996).

6 Choy, W. Y. *et al.* Distribution of molecular size within an unfolded state ensemble using small-angle X-ray scattering and pulse field gradient NMR techniques. *J Mol Biol* **316**, 101-112, doi:10.1006/jmbi.2001.5328 (2002).

7 Hofmann, H. *et al.* Polymer scaling laws of unfolded and intrinsically disordered proteins quantified with single-molecule spectroscopy. *Proc Natl Acad Sci U S A* **109**, 16155-16160, doi:10.1073/pnas.1207719109 (2012).

8 Sherman, E. & Haran, G. Coil-globule transition in the denatured state of a small protein. *Proc Natl Acad Sci U S A* **103**, 11539-11543, doi:10.1073/pnas.0601395103 (2006).

9 Kjaergaard, M. *et al.* Temperature-dependent structural changes in intrinsically disordered proteins: formation of alpha-helices or loss of polyproline II? *Protein Sci* **19**, 1555-1564, doi:10.1002/pro.435 (2010).

10 Arbesu, M. *et al.* The Unique Domain Forms a Fuzzy Intramolecular Complex in Src Family Kinases. *Structure* **25**, 630-640 e634, doi:10.1016/j.str.2017.02.011 (2017).

11 Mittag, T. *et al.* Structure/function implications in a dynamic complex of the intrinsically disordered Sic1 with the Cdc4 subunit of an SCF ubiquitin ligase. *Structure* **18**, 494-506, doi:10.1016/j.str.2010.01.020 (2010).

12 Wells, M. *et al.* Structure of tumor suppressor p53 and its intrinsically disordered N-terminal transactivation domain. *Proc Natl Acad Sci U S A* **105**, 5762-5767, doi:10.1073/pnas.0801353105 (2008).

13 Metrick, C. M., Koenigsberg, A. L. & Heldwein, E. E. Conserved Outer Tegument Component UL11 from Herpes Simplex Virus 1 Is an Intrinsically Disordered, RNA-Binding Protein. *mBio* **11**, doi:10.1128/mBio.00810-20 (2020).

14 Lens, Z. *et al.* Solution structure of the N-terminal transactivation domain of ERM modified by SUMO-1. *Biochem Biophys Res Commun* **399**, 104-110, doi:10.1016/j.bbrc.2010.07.049 (2010).

15 Riback, J. A. *et al.* Innovative scattering analysis shows that hydrophobic disordered proteins are expanded in water. *Science* **358**, 238-241, doi:10.1126/science.aan5774 (2017).

16 Flanagan, J. M., Kataoka, M., Shortle, D. & Engelman, D. M. Truncated staphylococcal nuclease is compact but disordered. *Proc Natl Acad Sci U S A* **89**, 748-752, doi:10.1073/pnas.89.2.748 (1992).

17 Nath, A. *et al.* The conformational ensembles of alpha-synuclein and tau: combining single-molecule FRET and simulations. *Biophys J* **103**, 1940-1949, doi:10.1016/j.bpj.2012.09.032 (2012).
18 Mylonas, E. *et al.* Domain conformation of tau protein studied by solution small-angle X-ray scattering. *Biochemistry* **47**, 10345-10353, doi:10.1021/bi800900d (2008).
